# Safety and efficacy of *C9ORF72*-repeat RNA nuclear export inhibition in amyotrophic lateral sclerosis

**DOI:** 10.1101/2021.04.12.438950

**Authors:** Lydia M. Castelli, Luisa Cutillo, Cleide Dos Santos Souza, Alvaro Sanchez-Martinez, Ilaria Granata, Monika A. Myszczynska, Paul R. Heath, Matthew R. Livesey, Ke Ning, Mimoun Azzouz, Pamela J. Shaw, Mario R. Guarracino, Alexander J. Whitworth, Laura Ferraiuolo, Marta Milo, Guillaume M. Hautbergue

## Abstract

**Background:** Loss of motor neurons in amyotrophic lateral sclerosis (ALS) leads to progressive paralysis and death. Dysregulation of thousands of RNA molecules with roles in multiple cellular pathways hinders the identification of ALS-causing alterations over downstream changes secondary to the neurodegenerative process. How many and which of these pathological gene expression changes require therapeutic normalisation remains a fundamental question.

**Methods:** Here, we investigated genome-wide RNA changes in C9ORF72-ALS patient-derived neurons and *Drosophila*, as well as upon neuroprotection taking advantage of our gene therapy approach which specifically inhibits the SRSF1-dependent nuclear export of pathological *C9ORF72*-repeat transcripts. This is a critical study to evaluate (i) the overall safety and efficacy of the partial depletion of SRSF1, a member of a protein family involved itself in gene expression, and (ii) a unique opportunity to identify neuroprotective RNA changes.

**Results:** Our study demonstrates that manipulation of 362 transcripts out of 2,257 pathological changes in C9ORF72-ALS patient-derived neurons is sufficient to confer neuroprotection upon partial depletion of SRSF1. In particular, expression of 90 disease-altered transcripts is fully reverted upon neuroprotection leading to the characterisation of a human C9ORF72-ALS disease-modifying gene expression signature. These findings were further investigated *in vivo* in diseased and neuroprotected *Drosophila* transcriptomes, highlighting a list of 21 neuroprotective changes conserved with 16 human orthologues in patient-derived neurons. We also functionally validated the high therapeutic potential of one of these disease-modifying transcripts, demonstrating that inhibition of ALS-upregulated human KCNN1-3 (*Drosophila* SK) voltage-gated potassium channel orthologs mitigates degeneration of human motor neurons as well as *Drosophila* motor deficits.

**Conclusions:** Strikingly, manipulating the expression levels of a small proportion of RNAs is sufficient to induce a therapeutic effect, further indicating that the SRSF1-targeted gene therapy approach is safe in the above preclinical models as it does not disrupt globally gene expression. The efficacy of this intervention is also validated at genome-wide level with therapeutically-induced RNA changes involved in the vast majority of biological processes affected in C9ORF72-ALS. Finally, the identification of a characteristic signature with key RNA changes modified in both the disease state and upon neuroprotection also provides potential new therapeutic targets and biomarkers.

## Background

Polymorphic GGGGCC hexanucleotide-repeat expansions in the *C9ORF72* gene cause the most common forms of familial amyotrophic lateral sclerosis (ALS) and frontotemporal dementia (FTD) [1, 2], a spectrum of fatal diseases which respectively lead to progressive death of motor neurons and of neurons in the frontal/temporal lobes of the brain. While ALS causes progressive paralysis and death usually within 2-5 years from symptom onset [3, 4], FTD patients present with altered cognitive features and psychological disinhibition [5]. No effective disease-modifying therapy is currently available for these diseases.

Alteration of multiple biological processes in ALS leads to pathophysiological changes including stress responses, mitochondrial dysfunction, alterations in axonal transport, autophagy and protein clearance, cell death, excitotoxicity, neuroinflammation and dysregulated astrocyte-motor neuron crosstalk among others [6] in association with widespread alteration of the RNA metabolism and gene expression [7]. Consistent with this, the expression levels, alternative splicing and poly-adenylation site usage of thousands of transcripts are altered in *C9ORF72* repeat-expansion carriers [8, 9]. This raises serious challenges for the identification of altered transcripts causing neurodegeneration since an unknown proportion of RNA changes may occur as a secondary downstream process resulting from disease progression and dysregulation of RNA-processing factors. *C9ORF72* repeat-expansions contribute to neuronal injury through three non-mutually exclusive mechanisms: haploinsufficiency, repeat-RNA sequestration of proteins and repeat-associated non-AUG (RAN) translation of neurotoxic dipeptide-repeat proteins (DPRs), the latter being considered one of the main drivers of neurodegeneration [10, 11].

We recently showed that repeat-RNA sequestration of the splicing factor and nuclear export adaptor SRSF1 (serine/arginine-rich splicing factor 1 previously known as ASF/SF2) triggers the nuclear export of intron-1-retaining *C9ORF72* repeat transcripts which lead to the cytoplasmic RAN translation of DPRs [12]. Conversely, depleting SRSF1 confers a selective disease-modifying therapeutic approach for neuroprotection in patient-derived neurons and *Drosophila* models of disease [12]. Interaction of dephosphorylated SRSF1 with NXF1 (nuclear export factor 1) upon completion of splicing [13, 14] induces handover of the mRNA through remodeling of NXF1 into a high RNA-binding conformation [15, 16] that licenses the nuclear export process via transient interactions of NXF1 with the protruding FG-repeats of nucleoporins which decorate the channel of the nuclear pore [17, 18]. The remodeling mechanism of NXF1 provides in turn a control mechanism to retain un-spliced transcripts within the nucleus [19, 20]. *C9ORF72* intron-1 repeat-RNA sequestration of SRSF1 is thought to trigger inappropriate remodeling of NXF1 into the high RNA affinity mode that promotes the nuclear export of pathological *C9ORF72* pre-mRNAs which retain the intron-1 repeat expansions [12, 21].

The genome-wide functions of SRSF1 have not yet been investigated in neurons. In proliferative cells, it contributes to shaping the transcriptome through several RNA-processing functions involved in: (i) constitutive and alternative splicing [22–24]; (ii) nuclear export of mRNAs [13,15,20,25]; (iii) RNA stability/ surveillance [26] and (iv) release of paused RNA polymerase II from promoters [27]. The functions of SRSF1 have been extensively studied in immortalised and cancer cells, where overexpression leads to altered splicing functions linked to transformation and oncogenesis [28, 29]. SRSF1 plays an essential role in the tissue-specific splicing of the CaMKIIδ (Ca^2+^/calmodulin-dependent kinase IIδ) transcript that occurs during embryonic development of the heart, while in contrast, SR-rich proteins are dispensable to the viability of mature cardiomyocytes [30]. Accordingly, individual depletion (>90%) of each of the conserved SRSF1-7 proteins only affected the SRSF1-dependent nuclear export of 225 transcripts in immortalised cells and approximately 100-400 transcripts for the other SRSF factors, indicating that the NXF1-dependent nuclear export adaptor function involves redundancy and/or cooperation [25]. Taken together, further investigation is required to understand the genome-wide contribution of SRSF1 function in a post-mitotic neuronal context.

In this study, we examined the genome-wide therapeutic mechanisms by which SRSF1 depletion confers neuroprotection through investigation of whole-cell and cytoplasmic transcriptomes from healthy control and C9ORF72-ALS *Drosophila* and patient-derived neurons. Strikingly, the nuclear export inhibition of *C9ORF72* repeat transcripts led to expression changes for ∼250 human mRNAs involved in multiple disease pathways out of ∼2,250 changes in C9ORF72-ALS patient-derived neurons, indicating that, while the neurodegenerative process is characterised by a large number of gene expression changes, a small proportion of transcript changes is sufficient to suppress the neurodegenerative process. The analysis of SRSF1-depleted C9ORF72-ALS *Drosophila* transcriptomes led to a similar observation with conserved manipulation of cellular pathways. Moreover, levels of approximately one third of SRSF1-RNAi induced neuroprotective changes are completely reversed compared to the disease state, providing small *in vitro* and *in vivo* disease-modifying gene expression signatures. Finally, based on the integration of human-*Drosophila* transcriptomes that identified 16 conserved human neuroprotective transcript changes, we used pharmacological inhibition and genetic manipulation to show that the expression levels of a conserved small conductance Ca^2+^-activated potassium channel, which is increased in the ALS models, can be manipulated to mitigate the death of human C9ORF72-ALS motor neurons as well as the neurodegeneration-associated locomotor deficits in *Drosophila*.

## Methods

### Experimental cell and animal models

#### Patient-derived iNPC lines and differentiation into iNeurons

The human iNeuron cells were derived from iNPC lines of three healthy controls and three C9ORF72-ALS patients (Table 1). They have been used in multiple studies including the manuscript describing the neuroprotective role of the SRSF1-RNAi depletion [12]. The following Research Resource Identifiers have been registered for human iNPC lines 154 (RRID:CVCL_XX76), 155 (RRID:CVCL_UF81), 3050 (RRID: CVCL_UH66), 78 (RRID:CVCL_UF8), 183 (RRID:CVCL_UF85) and 201 (RRID:CVCL_UF86). Control and patient iNeurons were differentiated and cultured exactly as described previously [12] in 6-well plates (200,000 cells per well) and transduced with 5MOI LV-Control-RNAi or LV-*SRSF1*-RNAi lentivirus for 5 days.

#### Patient derived iPSC lines and differentiation into motor neurons

The human motor neurons were derived from human induced pluripotent stem cells (iPSC) lines (Table 1). iPSC cells derived from 2 patients carrying the mutation in C9orf72 (CS28iALS-C9nxx; RRID:CVCL_W558 and CS29iALS-C9nxx; RRID:CVCL_W559) and 1 iPSC cell derived from an unaffected control (CS14iCTR-21nxx; RRID:CVCL_JK54) were obtained from Cedars-Sinai, a nonprofit academic healthcare organization. The cell line control MIFF1 (RRID:CVCL_1E69) [31] was kindly provided by Prof Peter Andrews and Dr Ivana Barbaric (Centre for Stem Cell Biology, The University of Sheffield). iPSCs were maintained in Matrigel-coated plates (Corning®) according to the manufacturer’s recommendations in complete mTeSR™-Plus™ Medium (StemCell Technologies). Cultures were replenished with fresh medium every day. Cells were passaged every four to six days as clumps using ReLeSR™ (StemCell Technologies) an enzyme-free reagent for dissociation according to the manufacturer’s recommendations. For all the experiments in this study, iPSCs were used between passage 20 and 35, all iPSCs were cultured in 5% O_2_, 5% CO_2_ at 37°C. For Motor Neuron differentiation, neural differentiation of iPSCs was performed using the modified version dual SMAD inhibition protocol [32]. Briefly iPS cells were transferred to Matrigel-coated plates. On the day after plating (day 1), after the cells have reached ∼100% confluence, the cells were washed once with PBS and then the medium was replaced with neural medium (50% of KnockOut™ DMEM/F-12 (ThermoFisher Scientific), 50 % of Neurobasal (ThermoFisher Scientific), 0.5× N2 supplement (ThermoFisher Scientific), 1x Gibco® GlutaMAX™ Supplement (ThermoFisher Scientific), 0.5x B-27 (ThermoFisher Scientific), 50 U/ml penicillin (Lonza) and 50 mg/ml streptomycin(Lonza), supplemented with SMAD inhibitors (DMH-1 2 µM; SB431542-10 µM and CHIR99021 3 µM [ [Tocris]). The medium was changed every day for 6 days. On day 7, the medium was replaced for neural medium supplemented with DMH-1 2 µM, SB431542-10 µM and CHIR 1 µM, All-Trans Retinoic Acid 0.1 µM (RA; StemCell Technologies) and Purmorphamine 0.5 µM (PMN; Tocris). The cells were kept in this medium until day 12 when is possible to see a uniform neuroepithelial sheet. The cells were then split 1:6 with Accutase (Gibco™) onto matrigel substrate in the presence of 10 µM of rock inhibitor (Y-27632 dihydrochloride; Tocris), giving rise to a sheet of neural progenitor cells (NPC). After 24 hours of incubation, the medium was changed to neural medium supplemented with RA 0.5 µM and PMN 0.1 µM, and changed every day for 6 more days. On day 19 the motor neuron progenitors were split with accutase onto matrigel-coated plates and the medium was replaced with neural medium supplemented with RA 0.5 µM, PMN 0.1 µM, compound E 0.1 µM (Cpd E; Tocris), BDNF 10ng/mL (ThermoFisher Scientific), CNTF 10ng/mL (ThermoFisher Scientific) and IGF 10ng/mL (ThermoFisher Scientific). The cells were then fed alternate days with neuronal medium until day 40.

#### Drosophila stocks and husbandry

Flies were raised under standard conditions in a humidified, temperature-controlled incubator with a 12 h: 12 h light:dark cycle at 25 °C, on food consisting of agar, cornmeal, molasses, propionic acid and yeast. Transgene expression was induced using the tissue specific D42-GAL4 or nSyb-GAL4 driver. The following strains were obtained from the Bloomington Drosophila Stock Center (RRIDs indicated): UAS-Luciferase RNAi (BDSC_31603), D42-GAL4 (BDSC_8816), nSyb-GAL4 (BDSC_51635), SK^Mi10378^ (BDSC_55475), SK^Mi09020^ (BDSC_51246). The following strains were obtained from the Vienna *Drosophila* Resource Center: UAS-SF2-RNAi (VDRC v27776). UAS-G4C2×3 and UAS-G4C2×36 lines [33] were a gift from Prof Adrian M. Issacs (Department of Neurodegenerative Disease, UCL Institute of Neurology, London). Unless otherwise stated, all experiments were conducted using male flies.

### Pharmacological treatments

In order to evaluate the neuroprotective potential of apamin (Sigma Aldrich, 178270), a potent antagonist of calcium-activated potassium channels KCNN1 and KCNN3 [34], motor neurons derived from iPSC cells from unaffected controls and C9ORF72-ALS patients were exposed to apamin (0.1-10 µM) diluted in neuronal medium for 72 hours. Cells treated with dimethyl sulfoxide (DMSO; Sigma Aldrich), the vehicle for dilution of apamin, were used as a control.

### Total, nuclear and cytoplasmic RNAs fractionation from patient-derived iNeurons

Three wells of a 6-well plate were lysed were lysed in Reporter lysis buffer (*Promega*) for 10 min on ice before centrifugation at 17,000 *g*, 5 min, 4 °C for subsequent western blot and total RNA extractions. Six wells of a 6-well plate were used for nuclear/cytoplasmic fractionations that were performed as described previously [12]. Briefly, cells were lifted from the plates in DEPC PBS, pelleted by centrifugation at 400 *g* for 5 min and quickly washed with hypotonic lysis buffer (10 mM HEPES pH 7.9, 1.5 mM MgCl_2_, 10 mM KCl, and 0.5 mM DTT). Cells were then lysed in hypotonic lysis buffer containing 0.16 U µl^−1^ Ribosafe RNase inhibitors (Bioline), 2 mM PMSF and SIGMAFAST Protease Inhibitor Cocktail tablets, EDTA free (Sigma-Aldrich) for 15 minutes on ice. All lysates underwent differential centrifugation (1,500 *g*, 3 min, 4 °C then 3,500 *g*, 8 min, 4 °C and then 17,000 *g*, 1 min, 4 °C) transferring the supernatants to fresh tubes after each centrifugation. The pellets from centrifugation at 1,500 *g* for 3 min were lysed in Reporter lysis buffer (Promega) containing 0.16 U µl^−1^ Ribosafe RNase inhibitors, 2 mM PMSF and Protease Inhibitors for 10 min on ice and subsequently centrifuged at 17,000 *g*, 5 min, 4 °C to obtain the nuclear fractions. RNA was extracted from total, nuclear and cytoplasmic fractions as detailed below. Equal volumes of total, nuclear and cytoplasmic lysates were resolved by SDS–PAGE, electroblotted onto nitrocellulose membrane and probed using mouse anti-SSRP1 [1/500 dilution; Abcam #ab26212] and chicken anti-class III b-tubulin (TUJ1) [1/500; Millipore #AB9354] antibodies. SSRP1 was detected using HRP-conjugated mouse secondary antibody [1/10000; Promega #W4021] and TUJ1 was detected using HRP-conjugated chicken secondary antibody [1/10000; Promega #G1351].

#### RNA extraction

250 µl iNeuron total, nuclear or cytoplasmic extracts were added to 750 µl PureZOL™ (Bio-Rad) to extract the RNA. Total *Drosophila* RNA was extracted from *Drosophila* heads which had been ground to a powder under liquid nitrogen before addition of 250 µl reporter lysis buffer (Promega) and addition of 750 µl PureZOL™ to the frozen lysates to extract the RNA. Briefly, lysates were cleared by centrifugation for 10 min at 12,000 *g* at 4°C. One fifth the volume of chloroform was added and tubes were vigorously shaken for 15 s. After 10 min incubation at room temperature, tubes were centrifuged 12,000 *g*, 10 min, 4 °C and supernatants collected. RNA was precipitated for 30 min at room temperature with an equal volume of isopropanol and 2 µl glycogen and subsequently pelleted at 12,000 *g,* 20 min, 4 °C. Pellets were washed with 70% DEPC ethanol and re-suspended in DEPC water. All PureZol™ extracted RNA samples were treated with DNaseI (Sigma Aldrich) and quantified using a Nanodrop (NanoDropTechnologies). Fractionated extracts were subjected to RNA extraction using Direct-zol™ RNA microprep kits (Zymo Research) following the manufacturer’s protocol, including the recommended in-column DNase I treatment and quantified using a Nanodrop. RNA quality was then assessed using a eukaryote total RNA Nano 6000 Kit (Agilent Technologies) prior to high depth RNA sequencing or microarray analysis and qRT-PCR.

### Quantitative RT-PCR (qRT-PCR)

Following quantification, 1 µg RNA (iNeurons samples) or 2 µg RNA (*Drosophila* samples) was converted to cDNA using BioScript Reverse Transcriptase (Bioline). qRT–PCR primers were designed using Origene or Primer-BLAST and validated using a 1 in 4 serial template dilution series (standard curve with *R*^2^>0.97). qRT–PCR reactions were performed in duplicate using the Brilliant III Ultra-Fast SYBR Green QPCR Master Mix (Agilent Technologies) on a C1000 Touch™ thermos Cycler CFX96™ Real-Time System (BioRAD) using an initial denaturation step, 45 cycles of amplification (95°C for 30 s; 60°C for 30 s; 72°C for 1 min) prior to recording melting curves. qRT–PCR data was analysed using CFX Manager™ software (Version 3.1) (BioRAD) and GraphPad Prism (Version 7). The sequences of qPCR primers are provided in Supplementary information.

### Immunofluorescence microscopy

For immunostaining, cells were washed with phosphate-buffered saline (PBS) and fixed with 4% paraformaldehyde (Sigma Aldrich) for 10 min at room temperature. After fixation samples were washed three times with PBS and permeabilized with 0.3% Triton X-100 diluted in PBS for 5 min. The cells were subsequently blocked in 5% Donkey serum (DS; Millipore) for 1 h. After blocking, cell cultures were incubated with the appropriate primary antibodies (rabbit anti-Nestin, 1:500 [Biolegend #841901]; mouse anti-Pax6, 1:200 [Millipore #MAB5552]; mouse anti-Tuj1, 1:1000 [Biolegend #801201]; mouse anti-NeuN, 1:1000 [Biolegend #SIG-39860]; goat anti-Chat, 1:100 [Millipore #AB144P]; mouse anti-SM132 [Biolegend #801701], 1:500; guinea pig anti-Map2 [Synaptic Systems #188004], 1:1000; rabbit anti-Caspase 3, 1:200 [Millipore #AB3623]) diluted in PBS containing 1% of DS overnight. Cells were then washed with PBS three times. Fluorescent secondary antibodies (Alexa Fluor 488, 555, 594 or 647, diluted 1:400 with DS [ThermoFisher Scientific #A-21202, #A-21432, A-21450, #A-32744, #A-21206]) were subsequently added to the cells and incubated for 1h. The samples were washed with PBS three more times and incubated with 1.0 mg/mL 4,6-diamidino-2-phenylindole (DAPI; Sigma Aldrich) for nuclear staining. As a specificity control, all experiments included cultures where the primary antibodies were not added. Non-specific staining was not observed in such negative control conditions. RNA foci were visualized using RNA fluorescence *in* situ hybridization (FISH) as described previously [35]. Images were taken with the Opera Phenix^™^ High Content Screening System at ×40 magnification using the Harmony™ Image analysis system. We used 405, 488 and 594 nm and 647 lasers, along with the appropriate excitation and emission filters. These settings were kept consistent while taking images from all cultures.

### High-content automated imaging microscopy

To investigate whether apamin protects motor neurons (MNs) from cells death, MNs were treated with apamin for 72h (0.1-10 µM) and then were stained for active caspase 3, a typical apoptotic marker. MNs were plated at 2×10^4^ cells per well on matrigel-coated 96-well plates. After treatment, the cells were fixed and stained for active caspase 3 and MAP2, which was used as a marker to define the boundary of cells and DAPI for nuclear staining. A quantitative imaging analysis of the MN was conducted through the The Opera Phenix^™^ High Content Screening System at × 40 magnification using the Harmony™ Image analysis system. The following morphological features were assessed for both treated and control: percentage caspase-3 positive cells and the number of fragmented nuclei. At least 25 fields were randomly selected and scanned per well of a 96-well plate in triplicate. To identify and remove any false readings generated by the system, three random treated and untreated wells were selected and counted manually (blinded to groups).

#### *Drosophila* locomotor and lifespan assays

The startle induced negative geotaxis (climbing) assay was performed using a counter-current apparatus. Briefly, 20–50 flies were placed into the first chamber, tapped to the bottom, and given 10 s to climb a 10 cm distance. This procedure was repeated five times (five chambers), and the number of flies that remained within each chamber counted. The weighted performance of several groups of flies for each genotype was normalized to the maximum possible score and expressed as Climbing index [36]. For larval crawling assays, wandering third instar larvae were placed in the centre of a 1% agar plate and left to acclimatise for 30 seconds, after which the number of peristalsis waves that occurred in the following minute were recorded.

### Bioinformatics analysis

#### Next generation RNA Sequencing (RNA-seq)

Total RNA samples with RNA integrity numbers (RIN) comprised between 9.3 and 10.0 were sent to the Centre of Genomic Research at the University of Liverpool for RNA-seq (project LIMS14705). Dual-indexed strand-specific RNA-seq library were prepared from the submitted total RNA samples using RiboZero rRNA depletion and the NEBNext Ultra Directional RNA library preparation kits (*New England Biolabs*). Paired-end 2×150bp sequencing was performed on an *Illumina HiSeq 4000* platform. RNA-seq reads were quality-checked and trimmed by the Centre of Genomic Research at the University of Liverpool. Fastaq files were then aligned to the Human Genome GRCh38 using STAR 2.7 aligner [37] and the ensembl built on a University of Sheffield Unix cluster. The RNA-seq data have been deposited in *Gene Expression Omnibus* (GEO) under accession number GSE139900. An average of 104 millions 150 bp paired-end sequencing reads were obtained (Table S1). Approximately 5-10% of the genome is stably transcribed in human cell lines [38, 39]. The size of the human genome is 3×10^9^ bp. Therefore according to the Lander/Waterman equation (Coverage = read length x number of reads / size of dataset): Transcriptome coverage = (2 x 150 bp x 104×10^6^ reads) / (3×10^9^ bp x 10/100) = 104 fold.

#### Comparison of transcription profiles from fibroblasts and fibroblast-derived cells

Microarray data previously obtained (GEO Accession number GSE87385) from human fibroblasts, induced astrocytes and induced oligodendrocytes were compared to the transcriptome of induced neurons and post-mortem laser captured motor neurons (GEO Accession number GSE29652). After merging all expression data based on Gene ID (Official Gene Symbol), we obtained 10,688 gene transcripts that were annotated on both the microarray platform used to define the transcriptome of fibroblasts, induced astrocytes, oligodendrocytes and post-mortem motor neurons, as well as the RNA-seq data relative to induced human neurons. Expression data of these 10,688 transcripts were normalised across platforms based on expression of housekeeping genes and groups visualised using the *Qlucore* visualisation software. Principal component analysis (PCA) plots were obtained by performing multi-group comparison analysis applying p-value < 0.01.

#### RNA-seq analysis: generation of differentially-expressed transcript lists

Over 80% of trimmed RNA-Seq reads provided by the Centre for Genomics Research at the University of Liverpool were aligned to the human GRCh38.79 genome build using STAR2.7 [37]. Table S1 provides numbers and proportion of aligned reads. Quantification of transcripts abundance was done using the aligned .bam files with RSEM, a method to accurately quantify transcripts from RNA-Seq data with or without a reference genome [40]. Transcripts counts were then analysed with EdgeR [41] to normalise the data and quantify differential expression. Data normalisation was performed using the relative log expression (RLE) implemented in EdgeR. After normalisation, transcripts abundance was filtered for low and no reads. We retained transcripts for which counts per million (cpm) were greater or equal than 2 in at least two samples. Transcripts sequenced in each group were compared to one another to evaluate transcriptome coverage across our experimental conditions. For each comparison, we evaluated the biological variation using the maximisation of the negative binomial dispersion using the empirical Bayes likelihood function, as implemented in EdgeR. Differentially-expressed transcript isoforms were computed for fold change FC>2 and p-value p<0.05, which were evaluated using the quasi-likelihood (QL) methods with empirical-Bayes Test in EdgeR. Differentially-expressed transcripts were annotated using BioMart [42] based on their Ensembl transcript Id to recover gene symbol, gene description and biotype.

#### RNA-seq analysis: splicing

For the splicing analysis, sequencing reads were aligned to the GRCh38.79 genome build through STAR two-pass mode (v2.5.4b) [37]. The gtf file was used as a guide during the first pass to find the superset of all novel splice junctions that were then used in the second-pass to improve the consistency of alignment and quantification across these spliced transcripts. The DEXSeq module of Bioconductor was used to identify differential exon usage [43]. We used the python scripts provided by the package to annotate the genome and to count the reads overlapping the exons. The significance thresholds for differential exon usage were set at a Benjamini–Hochberg false discovery rate of 5%.

#### Microarray analysis

Drosophila_2 gene expression arrays (*Affymetrix*) were used in this study. Total RNA samples were prepared according to the manufacturers’ protocol. Briefly, 200 ng of total RNA was converted into cDNA using an oligo(dT) which also carries the binding site for T7 RNA polymerase. Following first strand synthesis, residual RNA was degraded by addition of RNaseH and a double stranded cDNA molecule was generated using DNA Polymerase I and DNA ligase. These cDNA molecules were used as a substrate for the T7 RNA polymerase to produce multiple copies of antisense RNA using an IVT labelling system. The cRNA molecules produced incorporated biotin labelled ribonucleotides, which acted as a target for the subsequent detection of hybridization, using fluorescently labelled streptavidin. 12.5 µg of cRNA molecules were heat fragmented and applied to the GeneChips in a hybridization solution according to the Affymetrix protocol. Hybridization took place overnight in a rotating hybridization oven at 60 rpm, 45°C for 16 hours. The GeneChip arrays were washed using the ThermoFisher Fluidics Station. After washing and development of the fluorescent signal, the GeneChip arrays were scanned using GC30007 scanner. Gene level differential expression analysis was carried out using Transcriptome Analysis Console 3.1 (*Affymetrix*). The .CEL files were loaded into the software and sorted into the appropriate groups. Data were normalised using the RMA algorithm and the resultant grouped .CHP files compared for differential analysis. The software used a one way between subject ANOVA (unpaired) to calculate expression level differences with default values fold change FC>2 and p value p<0.05. The microarray data have been deposited in *Gene Expression Omnibus* (GEO) under accession number GSE138592.

### Statistical analysis of data

For qRT-PCR data, either one-way or two-way ANOVA (analysis of variance) with Tukey’s correction for multiple comparisons was used. For *D. melanogaster* climbing assays, a Kruskal– Wallis nonparametric test with Dunn’s post-hoc correction for multiple comparisons was used and data reported as mean ± 95% CI. For *D. melanogaster* crawling assays One-way ANOVA parametric with Bonferroni’s multiple comparison test was used and data reported as mean ± SEM. Data were plotted using GraphPad Prism 7. Significance is indicated as follows; NS: non-significant, *P*≥0.05; \**P*<0.05; \*\**P*<0.01; \*\*\**P*<0.001; \*\*\*\**P*<0.0001.

## Results

### Identifying transcriptomes from healthy and C9ORF72-ALS human-derived neurons

Depleting *SRSF1* mRNA levels by approximately 60% confers neuroprotection through inhibition of the nuclear export and RAN translation of pathological *C9ORF72*-repeat transcripts [12]. To investigate the genome-wide mechanisms by which SRSF1 depletion confers neuroprotection and evaluate the safety of this manipulation, we identified whole-cell and cytoplasmic transcriptomes to investigate RNA changes at expression, splicing and nuclear export levels. Neurons derived from induced neural progenitor cells reprogrammed from the fibroblasts of three different ALS patients harbouring C9ORF72-repeat expansion mutations (C9ORF72-ALS) and three healthy controls (**Table 1**) were treated with lentivirus expressing either scrambled control-RNAi (C-RNAi) or SRSF1-RNAi (ΔSRSF1) prior to nuclear and cytoplasmic fractionation in the same conditions and cell lines previously used to show that partial depletion of SRSF1 confers neuroprotection [12]. Western blot analysis using antibodies directed against the SSRP1 chromatin remodelling factor confirmed the absence of nuclear contamination in the cytoplasmic fractions (Figure 1A). Similarly to our previous study, qRT-PCR quantification showed ∼60% depletion of SRSF1 mRNA levels upon treatment with SRSF1-RNAi lentivirus (Figure 1B). We also confirmed that the depletion of SRSF1 did not affect the splicing of *C9ORF72* intron-1 (Figure 1C) and specifically inhibited the nuclear export of *C9ORF72* transcripts retaining the pathological repeat-expansions in intron-1, showing nuclear accumulation and concomitant cytoplasmic decrease of these repeat transcripts (Figure 1D).

**Fig. 1.**
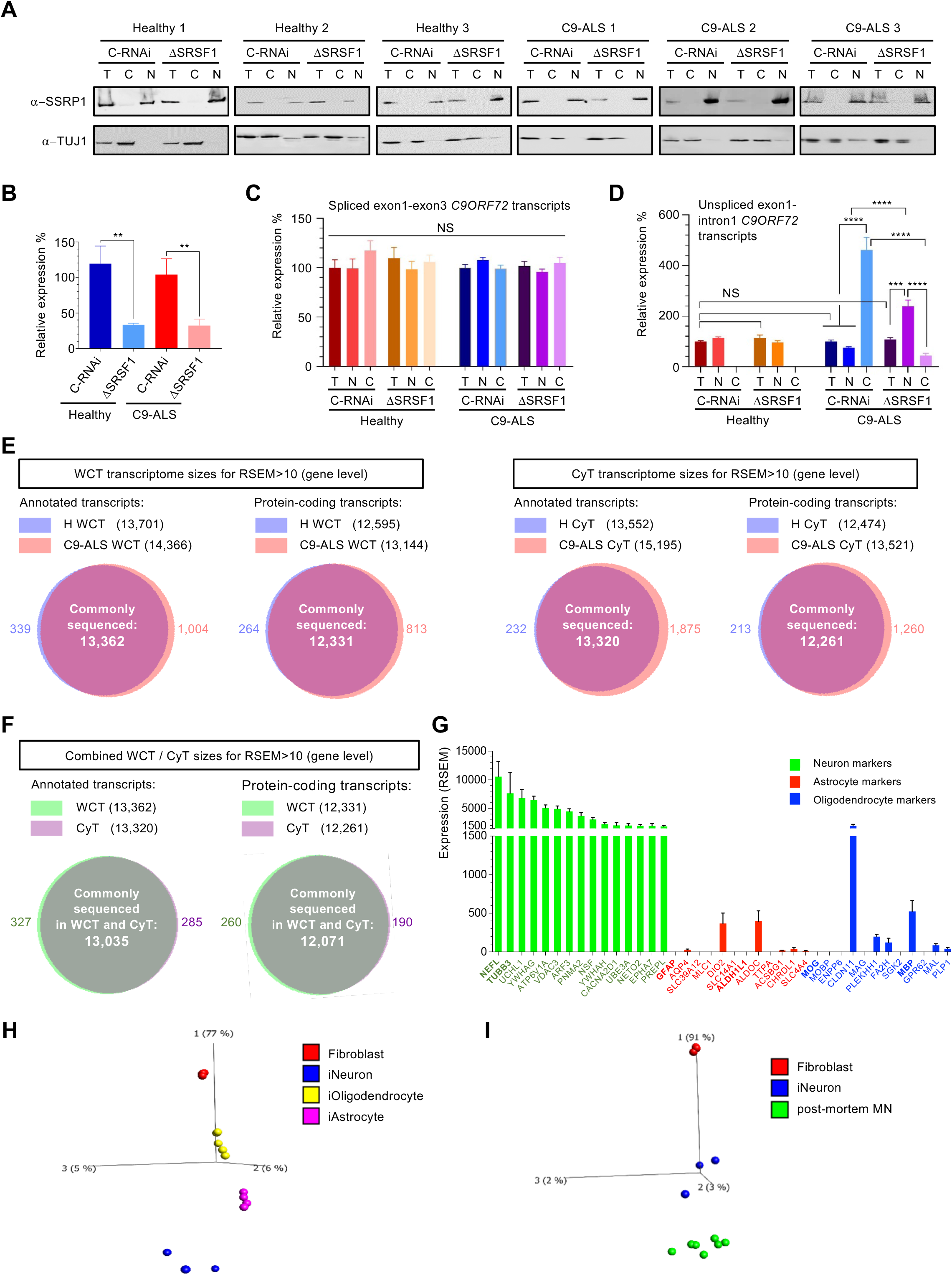
Generation of whole-cell and cytoplasmic transcriptomes from healthy and C9ORF72-ALS patient-derived neurons. (**A**) Three healthy control and three C9ORF72-ALS (C9-ALS) lines of patient-derived neurons were treated with Ctrl-RNAi (C-RNAi) or SRSF1-RNAi (ΔSRSF1) prior to whole-cell (T) lysis or nuclear (N) and cytoplasmic (C) fractionation. Western blots were probed for the nuclear chromatin remodelling SSRP1 factor and the neuronal cytoplasmic marker TUJ1. (**B**) Relative expression levels of *SRSF1* mRNA in whole-cell patient-derived neurons prepared in A were quantified using qRT-PCR in biological triplicates following normalization to U1 snRNA levels and to 100% for healthy neurons treated with C-RNAi (mean ± SEM; one-way ANOVA with Tukey’s correction for multiple comparisons, **: p<0.01; N (qRT-PCR reactions)=3). (**C**) Total, nuclear and cytoplasmic levels of intron1-spliced C9ORF72 transcripts (as measured by the exon1-exon3 junction) were quantified using qRT-PCR in biological triplicates following normalization to U1 snRNA levels and to 100% for whole-cell healthy neurons treated with C-RNAi (mean ± SEM; one-way ANOVA with Tukey’s correction for multiple comparisons, NS: non-significant; N (qRT-PCR reactions)=3). (**D**) Total, nuclear and cytoplasmic levels of unspliced C9ORF72 transcripts retaining intron1 (as measured by the exon1-intron1 junction) were quantified using qRT-PCR in biological triplicates following normalization to U1 snRNA levels and to 100% for whole-cell healthy neurons treated with C-RNAi (mean ± SEM; one-way ANOVA with Tukey’s correction for multiple comparisons, NS: non-significant; ***: p<0.001; ****: p<0.0001; N (qRT-PCR reactions)=3). (**E**) Whole cell transcriptome (WCT; left) and cytoplasmic transcriptome (CyT, right) sizes for quantified transcripts at gene level. (**F**) Combined WCT and CyT sizes for quantified transcripts at gene level. (**G**) Abundance of neuron (green), astrocyte (red) and oligodendrocyte (blue) markers in the human derived neurons (combined from all 24 protein-coding transcriptomes shown in Fig. 1F). (**H**) Principal component analysis (PCA) plot representing transcriptomes from patient-derived neurons, astrocytes, oligodendrocytes and fibroblasts of origin. (**I**) Principal component analysis (PCA) plot representing transcriptomes from patient-derived neurons, human post-mortem motor neurons (MN) and fibroblasts of origin.

We next proceeded to extract total RNA from the whole-cell and cytoplasmic fractions to analyse genome-wide RNA expression changes using next generation RNA sequencing (RNA-seq). Independent triplicates of rRNA-depleted RNA sequencing libraries were subjected to high depth RNA sequencing (averaging 104 million reads per sample, >100-fold transcriptome coverage; **Table S1**) to investigate differential expression and splicing with high confidence in unrelated patient samples, which present a high level of genetic variability (Material and Methods). The lists of over 40,000 quantified transcript isoforms are presented in **Table S2** under 4 tabs corresponding to each of whole-cell transcriptomes (WCT) and cytoplasmic transcriptomes (CyT) for either healthy control (H) or C9ORF72-ALS (C9) patient-derived neurons treated with C-RNAi or SRSF1-RNAi lentivirus. Over 13,000 annotated transcripts and 12,000 protein-coding mRNAs were commonly sequenced at gene level across all conditions reflecting the generation of datasets with high transcriptome coverage without notable RNA sequencing bias between transcriptomes (Figure 1E-F). Based on cell type-specific transcriptome databases from human-derived brain cells [44] and mouse brains [45], we identified the top 20-expressed transcripts in our datasets that are present within (i) the top 1000-specific human-derived neurons and 2034-enriched mouse brain neurons; (ii) the top 354-specific human-derived astrocytes and 2616-enriched mouse brain astrocytes and (iii) the top 260-specific human-derived oligodendrocytes and 2227-enriched mouse brain oligodendrocytes (**Table** S**3**). This analysis showed that our differentiation protocol successfully yields cells with high expression of neuronal markers (such as *NEFL* and *TUBB3/TUJ1*) and low expression levels of transcripts known to be specifically enriched in astrocytes (such as *GFAP* or *ALDH1L1*) and oligodendrocytes (such as *MOG* or *MBP*) (Figure 1G). Moreover, a principal component analysis (PCA) comparing induced patient-derived neurons, astrocytes and oligodendrocytes to their parental fibroblasts clearly showed specific segregation of the 4 different cell types away from each other and from the fibroblasts of origin; with glial cells, i.e. astrocytes and oligodendrocytes clustering closer to each other than to neurons (Figure 1H). Moreover, our protocol of differentiation leads to patient-derived neuron transcriptomes which cluster close to human post-mortem motor neurons on the main component of a PCA plot comparing fibroblasts to motor neurons (Figure 1I). Overall, high depth RNA-seq transcriptomes have been generated from cells successfully differentiated into patient-derived neurons.

### Therapeutic depletion of SRSF1 preserves the transcriptomes of human-derived neurons

A multidimensional scale analysis of the 24 identified transcriptomes (whole-cell and cytoplasmic triplicates from healthy and C9ORF72-ALS neurons treated with control-or SRSF1-RNAi lentivirus) shows that the partial depletion of SRSF1 does not overall disrupt the transcriptomes, with maintenance of the genetic variability between individuals (Figure 2A). Differentially-expressed transcript isoforms were selected for fold change FC>2 and p-value p<0.05. Despite a marked reduction in the abundance of sequenced SRSF1 isoforms in the matched pair of neurons treated with control-or SRSF1-RNAi, the differential expression of SRSF1 transcripts, which was experimentally validated (Figure 1B), provided p-values overall not statistically significant due to the high variability of SRSF1 expression between individual subjects (2 to 5 fold; **Table S4**).

**Fig. 2.**
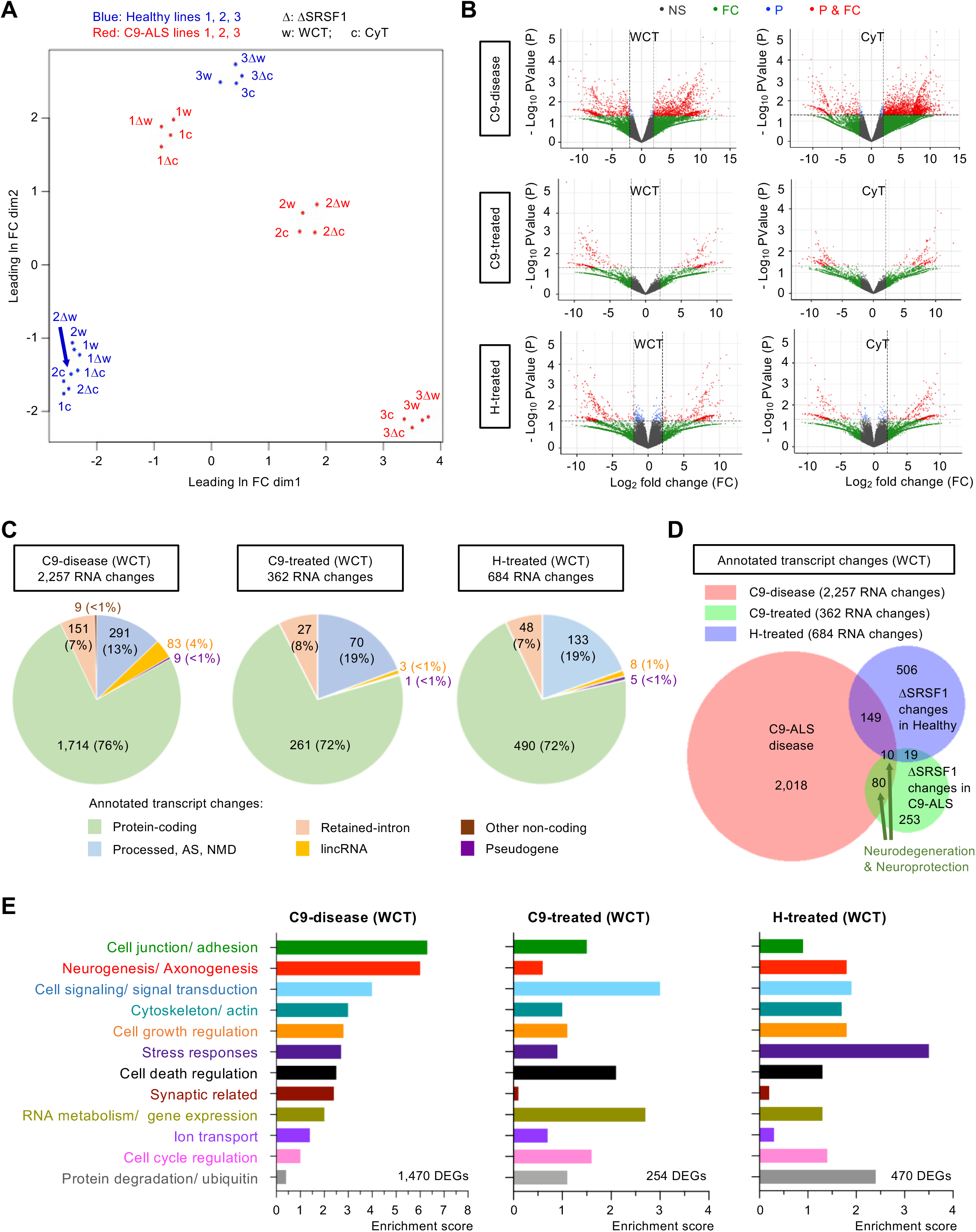
Genome-wide effects of the SRSF1 depletion in human healthy and C9ORF72-ALS neurons. (**A**) Multidimensional scale analysis of the WCT and CyT transcriptomes for healthy (blue) and C9-ALS (red) lines of patient-derived neurons treated with C-RNAi or SRSF1-RNAi (Δ). (**B**) Volcano plots representing the genome-wide distribution of differentially-expressed transcripts according to p-values and fold changes for C9-disease (H_C-RNAi versus C9_C-RNAi), C9-treated (C9_C-RNAi versus C9_SRSF1-RNAi) and H-treated (H_C-RNAi versus H_SRSF1-RNAi) neuron groups. NS: non-significant; FC: fold changes <2; P: p-values >0.05; P & FC: significant p-values (<0.05) & fold changes (>2). (**C, D**) Venn diagrams representing differentially-expressed transcripts at WCT level for C9-disease, C9-treated and H-treated groups. (**E**) Bar charts representing enrichment scores for the 12 top biological processes identified via functional annotation clustering for the C9-disease, C9-treated and H-treated groups. The therapeutic depletion of SRSF1 mitigates most of the disease-altered pathways.

Lists of differentially expressed transcript isoforms and differentially expressed genes (DEGs) are provided in **Table S5** for annotated and protein-coding transcripts in WCT and CyT transcriptomes. Changes were investigated in: (i) C9ORF72-ALS disease (H_C-RNAi versus C9_C-RNAi; tabs 1 & 4); C9ORF72-ALS neurons treated with SRSF1-RNAi (C9_C-RNAi versus C9_SRSF1-RNAi; tabs 2 & 5); healthy control neurons treated with SRSF1-RNAi (H_C-RNAi versus H_SRSF1-RNAi; tabs 3 & 6). Whole-cell expression levels of 2,257 transcripts (1,804 DEGs) are altered in C9ORF72-ALS versus healthy neurons (C9-disease group) while the therapeutic depletion of SRSF1 leads to the modulation of only 362 transcripts (351 DEGs; <1% of the transcriptome comprising transcripts from ∼42,500 coding and non-coding genes) in C9ORF72-ALS neurons treated with SRSF1-RNAi (C9-treated) and 684 transcript changes (642 DEGs; ∼1.5% transcriptome) in healthy neurons treated with SRSF1-RNAi (H-treated) (Figure 2B-C). The very small proportion of quantified transcript changes is in full agreement with the multidimensional scale analysis that did not show global alteration of SRSF1-depleted transcriptomes. Consistent with the mRNA splicing and nuclear export functions of SRSF1, 72% of manipulated RNAs in SRSF1-RNAi-treated neurons are protein coding transcripts (Figure 2C). However, the SRSF1-RNAi-induced transcript changes in healthy and C9ORF72-ALS neurons do not overlap (Figure 2D) reinforcing the concept that healthy control and C9-ALS transcriptomes are very diverse at a global level due to widespread alteration of RNA metabolism in the C9ORF72-ALS disease state (Figure 2A).

### Therapeutic manipulation of SRSF1 ameliorates multiple dysregulated pathways in C9ORF72-ALS neurons

A gene ontology (GO) analysis was performed with the protein-coding lists of WCT DEGs provided in table S5 using *DAVID 6.8* [46, 47]. The results are presented in **Table S6** under 3 tabs for the investigation of biological processes modulated in C9-disease, C9-treated and H-treated groups. Transcripts altered in C9-disease encode proteins involved in cell junction/ adhesion, neurogenesis/ axonogenesis, cytoskeleton, cell signaling, regulation of cell growth and cell death, stress responses, synaptic signaling, RNA metabolism/ gene expression, ion transport and cell cycle regulation (Figure 2E, C9-disease). Interestingly, despite the fact that the depletion of SRSF1 in healthy control and C9ORF72-ALS neurons led to different transcript changes (Figure 2D), the same pathways are manipulated upon SRSF1 depletion (Figure 2E, C9-treated versus H-treated). In agreement with the proliferative and oncogenic functions of SRSF1 [28, 29], the depletion of SRSF1 in healthy neurons leads to down-regulation of markers of the G1/S cell cycle transition and mitosis (Supplementary Figure S1A) as well as of cell proliferation and cancer markers [48, 49] (Supplementary Figure S1B). Interestingly, reducing the expression level of SRSF1 also promotes expression of transcripts involved in neuron differentiation, axonogenesis and synaptic transmission (Supplementary Figure S1C), suggesting additional roles of SRSF1 that may provide neuroprotective benefits beyond inhibiting the nuclear export of *C9ORF72* repeat transcripts and the production of DPRs.

As shown above and reported in the literature, a broad range of cellular processes are affected in C9ORF72-ALS. Remarkably, the neuroprotection conferred by the depletion of SRSF1 appears to act upon the vast majority of biological pathways dysregulated in disease with the exception of synaptic-related signaling which is poorly manipulated and protein degradation which is up-regulated upon neuroprotection. (Figure 2E, C9-disease versus C9-treated). Out of 2,257 RNA changes which occur in 1,804 genes in C9ORF72-ALS, manipulating the expression levels of 362 transcripts only (∼16%, including 261 protein-coding genes) is sufficient to confer neuroprotection *in vitro* (Figure 2C) suggesting that the large majority of RNA alterations are not directly related to pathogenesis but are likely to be downstream consequences of the neurodegenerative process. Importantly, this shows that therapeutic efficacy can be achieved without mitigating most of the disease-altered transcript changes. While the precise gene expression changes causing neurodegeneration still remain to be determined, the identification of 261 neuroprotective mRNAs offers new promising perspectives to understand disease pathophysiology and identify novel therapeutic targets for C9ORF72-ALS.

### Depletion of SRSF1 confers neuroprotection independently of genome-wide modulation of splicing and mRNA nuclear export

The genome wide effects of the therapeutic depletion of SRSF1 were next investigated at splicing and mRNA nuclear export levels. For the splicing analysis, exon reads were extracted from bam files and differences in exon usage were computed for C9-disease, C9-treated and H-treated neurons (Methods). This methodology allows the detection of differential splicing in an exon-centric manner, through the analysis of differential exon usage between the conditions under study, directly related to the diversity of genes and, hence, to alternative splicing. This analysis is independent of the knowledge of the exact transcript isoform(s) that can often generate misleading results since one exon can be shared between several assembled transcripts [50–52]. **Table S7** reports all statistically differentially expressed exons identified at a false discovery rate of 5% under 6 tabs for either WCT or CyT: C9-disease (tabs 1 & 4), C9-treated (tabs 2 & 5) and H-treated (tabs 3 & 6). **Table S8** provides a summary of the changes. Cytoplasmic exon usage changes are less noisy than the whole-cell samples that contain processing pre-mRNAs. Altered transcript isoforms that have been exported into the cytoplasm are also more likely to have functional consequences at the protein level. 99 differentially expressed cytoplasmic exons were identified in 77 genes in C9-disease, while no splicing changes were detected in neuroprotected C9ORF72-ALS neurons and only 6 were found in SRSF1-depleted healthy control neurons (Figure 3A). In particular, no changes were detected in the splicing of the *C9ORF72* transcripts in either C9ORF72-ALS or healthy control neurons in agreement with qRT-PCR assays in Figure 1C. These data indicate that the partial depletion of SRSF1, although neuroprotective, has no significant effect on the genome-wide splicing of transcripts.

**Fig. 3.**
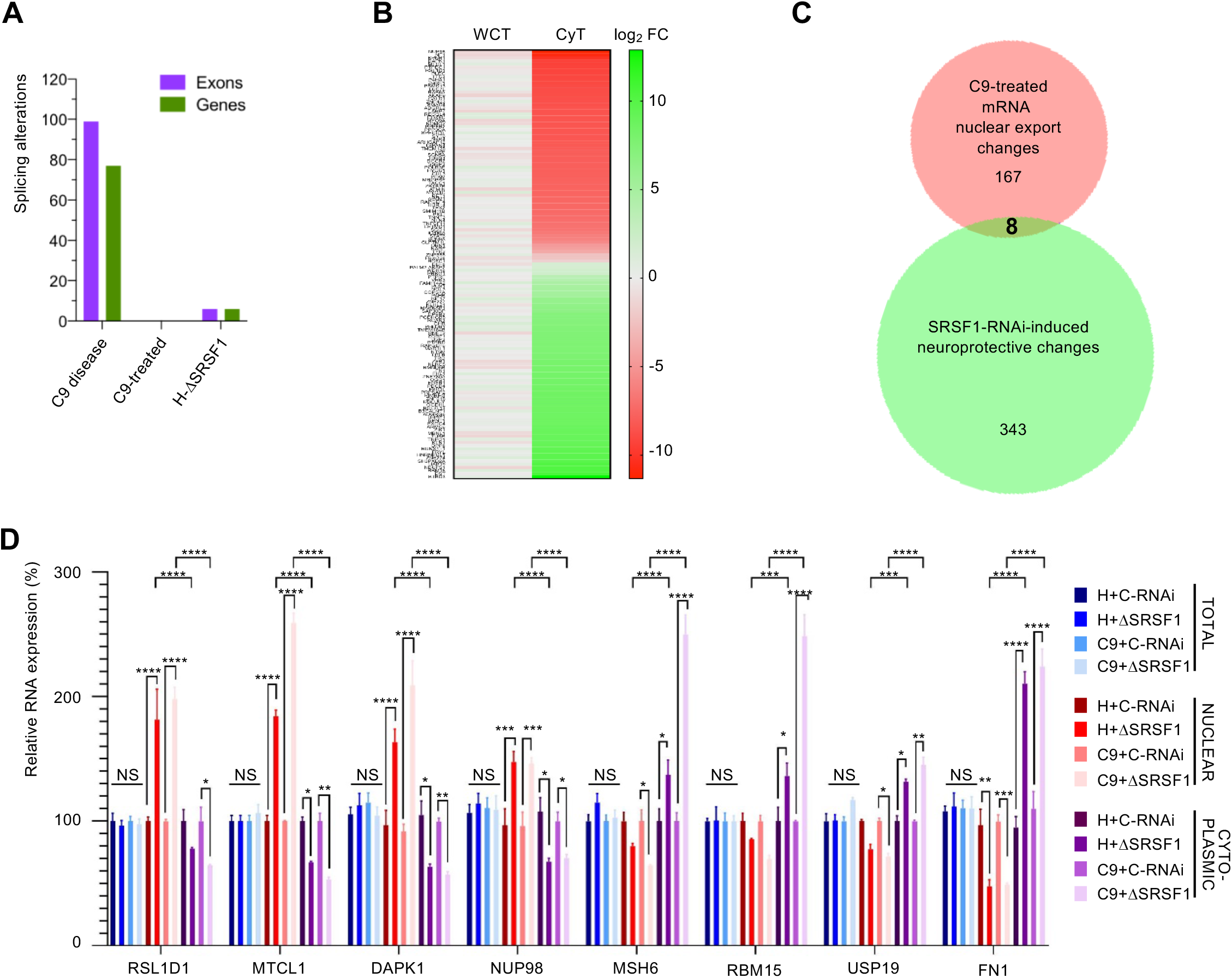
Genome-wide investigation of splicing and nuclear export in human neurons therapeutically depleted of SRSF1. (**A**) Bar chart representing the genome-wide number of identified splicing alterations at exon (purple) or gene (green) level for the C9-ALS-disease, C9-ALS-treated and SRSF1-depleted healthy neurons. (**B**) Genome wide nuclear RNA export analysis of the SRSF1 depletion in C9-ALS patient derived neurons. The heatmap represents transcript fold changes for FC>3 in WCT and FC>3 in CyT. Red labels shows down-regulated transcripts while green depicts upregulated transcripts. (**C**) Venn diagram representing the lists of RNA nuclear export alterations and SRSF1-RNAi-induced neuroprotective changes in C9-ALS neurons. (**D**) Relative RNA expression levels of *RSL1D1*, *MTCL1*, *DAPK1*, *NUP98*, *MSH6*, *RBM15*, *USP19* and *FN1* transcripts in total, nuclear and cytoplasmic fractions were quantified using qRT-PCR in biological triplicates following normalization to *U1* snRNA levels and to 100% for whole-cell healthy neurons treated with C-RNAi (mean ± SEM; two-way ANOVA with Tukey’s correction for multiple comparisons, NS: not significant; *: p<0.05, **: p<0.01, ***: p<0.001; ****: p<0.0001; N (qRT-PCR reactions)=3).

We also investigated the potential impact of partial depletion of SRSF1 on the genome-wide nuclear export of transcripts. Transcript abundance with FC >3 in the cytoplasmic transcriptomes were intersected with transcripts not significantly changing in the whole cell transcriptomes (FC <3) in a similar approach used for measuring the mRNA nuclear export dependence of SRSF1-7 proteins in a murine cell line [25]. The nuclear export of 177 annotated transcripts, which include 137 mRNAs, is altered in C9ORF72-ALS neurons treated with SRSF1-RNAi representing 0.4% of the transcriptome while 202 mRNAs have altered nuclear export in healthy neurons depleted of SRSF1 (**Table S9**). Approximately half of the transcripts either showed nuclear export inhibition or stimulation (Fig 3B; extended heat map with transcript IDs in Supplementary Figure S2), in full agreement with a previous report which identified nuclear export alterations of 225 transcripts upon depletion of over 90% of SRSF1 in a proliferating cell context [25]. The absence of overlap between transcripts with altered RNA nuclear export and the list of SRSF1-RNAi-induced neuroprotective changes (Figure 3C) suggests that neuroprotection is conferred independently of the nuclear export modulation of cellular transcripts. Importantly, partial depletion of SRSF1 does not alter the CaMKIIδ transcript essential to the developing heart [30] at expression, splicing or nuclear export level.

To experimentally validate the outputs of the bioinformatics analysis, we investigated some of the 4 most down-regulated and 4 most up-regulated transcripts of the altered nuclear export list (**Table S9**; tab C9-treated ann. transcripts) which represents a compilation of predicted data from both whole-cell and cytoplasmic transcriptomes. Using qRT-PCR assays, we showed that the expression levels of *RSL1D1*, *MTCL1*, *DAPK1*, *NUP98* and *MSH6*, *RBM15*, *USP19*, *FN1* are indeed respectively decreased and increased in the cytoplasm while the total levels remain unchanged and the nuclear expression levels are altered in the opposite direction (Figure 3D). Interestingly, fold changes for transcripts with the highest altered nuclear export were no more than 2.5 fold, indicating that the genome-wide effects of the SRSF1 depletion on nuclear export is limited (both in the numbers and FC of affected transcripts) reminiscent of the study which showed that the family of SRSF1-7 proteins play redundant and/or cooperative roles in the NXF1-dependent nuclear export adaptor function [25].

### Therapeutic manipulation of SRSF1 modulates multiple dysregulated pathways in C9ORF72-ALS *Drosophila*

Partial depletion of SRSF1 suppressed the *C9ORF72*-repeat neurodegeneration-associated locomotor deficits of G4C2×36 *Drosophila* [53] through the transgenic expression of *SRSF1*-specific RNAi sequences [12]. Here, we investigated the transcriptomes of *Drosophila* heads from the same lines: (i) healthy control flies expressing 3 G4C2 repeats and a luciferase-RNAi control (G4C2×3_C-RNAi); (ii) C9ORF72-ALS model expressing 36 G4C2 repeats and the RNAi control (G4C2×36_C-RNAi); (iii) C9ORF72-ALS-neuroprotected flies expressing 36 G4C2 repeats and the disrupted SRSF1 allele (G4C2×36_SRSF1-RNAi).

Lists of differentially expressed transcripts were identified for FC>2 and p<0.05 (Methods). **Table S10** reports changes in: (i) C9-disease (tab 1; G4C2×3_C-RNAi versus G4C2×36_C-RNAi), (ii) C9-treated (tab 2; G4C2×36_C-RNAi versus G4C2×36_SRSF1-RNAi), (iii) H vs C9-treated (tab 3; G4C2×3_C-RNAi versus G4C2×36_SRSF1-RNAi). 644 DEGs were identified in C9-disease flies while expression of 1,468 and 1,559 genes was respectively changed in the C9-treated and H vs C9-treated groups. Venn diagrams show that SRSF1-RNAi-induced therapeutic efficacy is achieved without normalizing all of the transcripts that are altered in disease and that expression levels of 346 transcripts are altered both in disease and upon neuroprotection (Figure 4A). The *DAVID* ontology analysis is available for all groups in **Table S11** (tabs 1, 2, 3). Transcripts altered in the C9ORF72-ALS *Drosophila* heads include biological processes previously identified in patient-derived neurons (such as cell signalling, stress responses, and ion transport) but also changes in defence/ immune response, lipid/ carbohydrate metabolisms, neurological process and learning/ memory (Figure 4B, C9-disease). On the other hand, the neuroprotective effects conferred by the depletion of SRSF1 act again remarkably upon the vast majority of pathways dysregulated in disease (Figure 4B, C9-treated). Interestingly, other cellular processes that were found altered in both C9-disease and C9-treated patient-derived neurons (such as protein degradation, gene expression, synaptic-related, cell death regulation, cytoskeleton and cell junction/adhesion) are also enriched in neuroprotected *Drosophila* heads (Figure 4B, C9-treated). This indicates that the inhibition of the RAN translation of C9ORF72-repeat transcripts mediated by the depletion of SRSF1 confers neuroprotection by inducing changes in conserved cellular pathways. This is amplified in the H vs C9-treated group which highlights the manipulated pathways that have been modified in C9-treated flies compared to the healthy animals and which confer neuroprotection in C9-ALS flies (Figure 4B). Highly enriched changes include transcripts involved in cell signaling, synaptic-related processes, neurogenesis/ axonogenesis and locomotion. Overall, the modulation of these biological processes appear particularly relevant to a *Drosophila* model of C9ORF72-ALS and the mitigation of the neurodegeneration associated locomotor deficits. Using qRT-PCR assays, we further experimentally validated that thioredoxin (*DHD*), the most downregulated transcript in C9ORF72-ALS *Drosophila*, and the sodium-dependent nutrient aminoacid transporter *NAAT1* in the ion transport pathway, exhibit reduced mRNA expression levels in disease while they are upregulated to normal levels upon SRSF1 depletion (Figure 4C) in agreement with the computed DEG lists (**Table S10**, C9-disease and C9-treated).

**Fig. 4.**
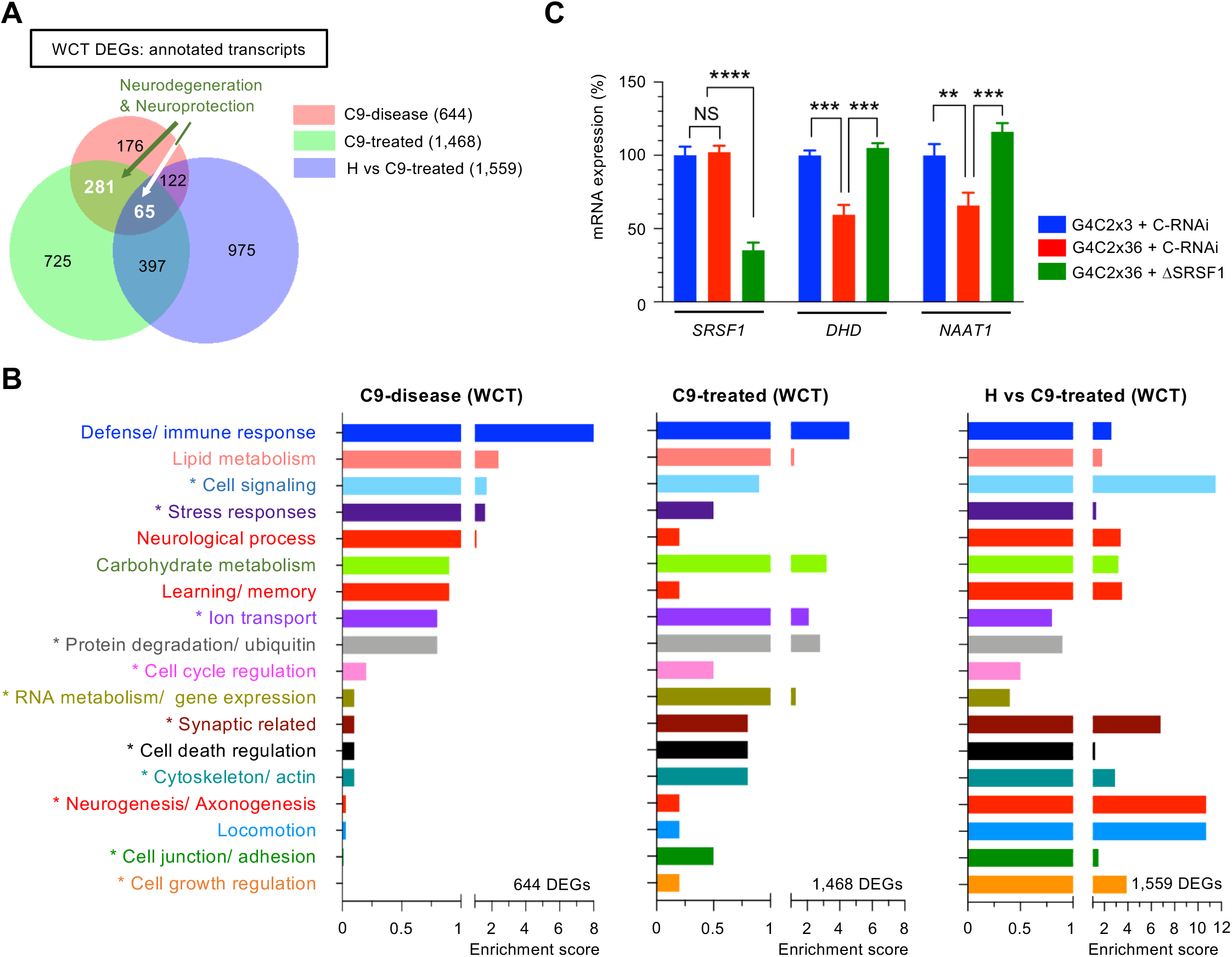
Genome-wide investigation of the partial depletion of SRSF1 in C9-disease and C9-treated *Drosophila* heads. (**A**) Venn diagram representing the numbers of annotated transcripts at gene level identified in the C9-disease (G4C2×3 + Ctrl-RNAi versus G4C2×36 + Ctrl-RNAi), C9-treated (G4C2×36 + Ctrl-RNAi versus G4C2×36 + SRSF1-RNAi) and H vs C9-treated (G4C2×3 + Ctrl-RNAi versus G4C2×36 + SRSF1-RNAi) groups. (**B**) Bar charts representing enrichment scores for biological processes identified via functional annotation clustering of the whole cell transcriptomes for the C9-disease, C9-treated and H vs C9-treated groups. Modulated pathways annotated with an asterisk (*) were also identified in the human patient-derived neuron transcriptomes. The therapeutic depletion of SRSF1 mitigates most of the disease-altered pathways. (**C**) Relative expression levels of *SRSF1*, *DHD* and *NAAT1* mRNAs were quantified using qRT-PCR for the indicated *Drosophila* lines in biological triplicates following normalization to Tub84b mRNA levels and to 100% for G4C2×3 + C-RNAi *Drosophila* heads (mean ± SEM; one-way ANOVA with Tukey’s correction for multiple comparisons; **: p<0.01, ***: p<0.001, ****: p<0.0001; N (qRT-PCR reactions)=3).

### Identifying conserved disease-modifying neuroprotective changes

The therapeutic depletion of SRSF1 provides a unique opportunity to investigate manipulated disease-modifying mechanisms through the identification of gene expression changes that occur in both C9ORF72-ALS and upon neuroprotection. The expression levels of only 90 transcripts affected in disease (∼4% out of 2,257 RNA changes) are also altered following SRSF1 depletion (Figure 5A). A clustered heat map represents how transcripts are modulated in diseased and neuroprotected neurons for each individual subject (Supplementary Figure S3). It is noteworthy that, despite genetic variability between individuals, the depletion of SRSF1 reverses altered expression levels for almost all disease-associated transcripts across the cases. This analysis was further validated by investigating the fold change values calculated for each of the 90 transcripts which have modified expression levels in the C9-disease and C9-treated groups (Figure 5B, enlarged heatmap in Supplementary Figure S4, transcript IDs and FC values in **Table S12 tab 1**). Remarkably, the altered expression levels of 88 transcripts, which include 68 mRNAs, are completely reversed upon SRSF1 depletion. We next sought to perform the same analysis using the *Drosophila* transcriptomes and investigated the 346 transcript changes that are both affected in disease and manipulated upon SRSF1-RNAi-dependent neuroprotection (Figure 5C) showing again, as for the human-derived neuron data, that the transcripts which are downregulated or upregulated in C9-disease are largely completely reversed in the neuroprotected C9-treated flies (Figure 5D, transcript IDs and FC values in **Table S12 tab 2**). The identification of disease-modifying gene expression signatures with transcripts involved in different biological functions that show reversed expression levels upon neuroprotection in both *in vitro* and *in vivo* models is entirely consistent with our previous conclusions indicating that the therapeutic manipulation of SRSF1 counteracts multiple disease-altered biological processes (Figure 2E and 4B).

**Fig. 5.**
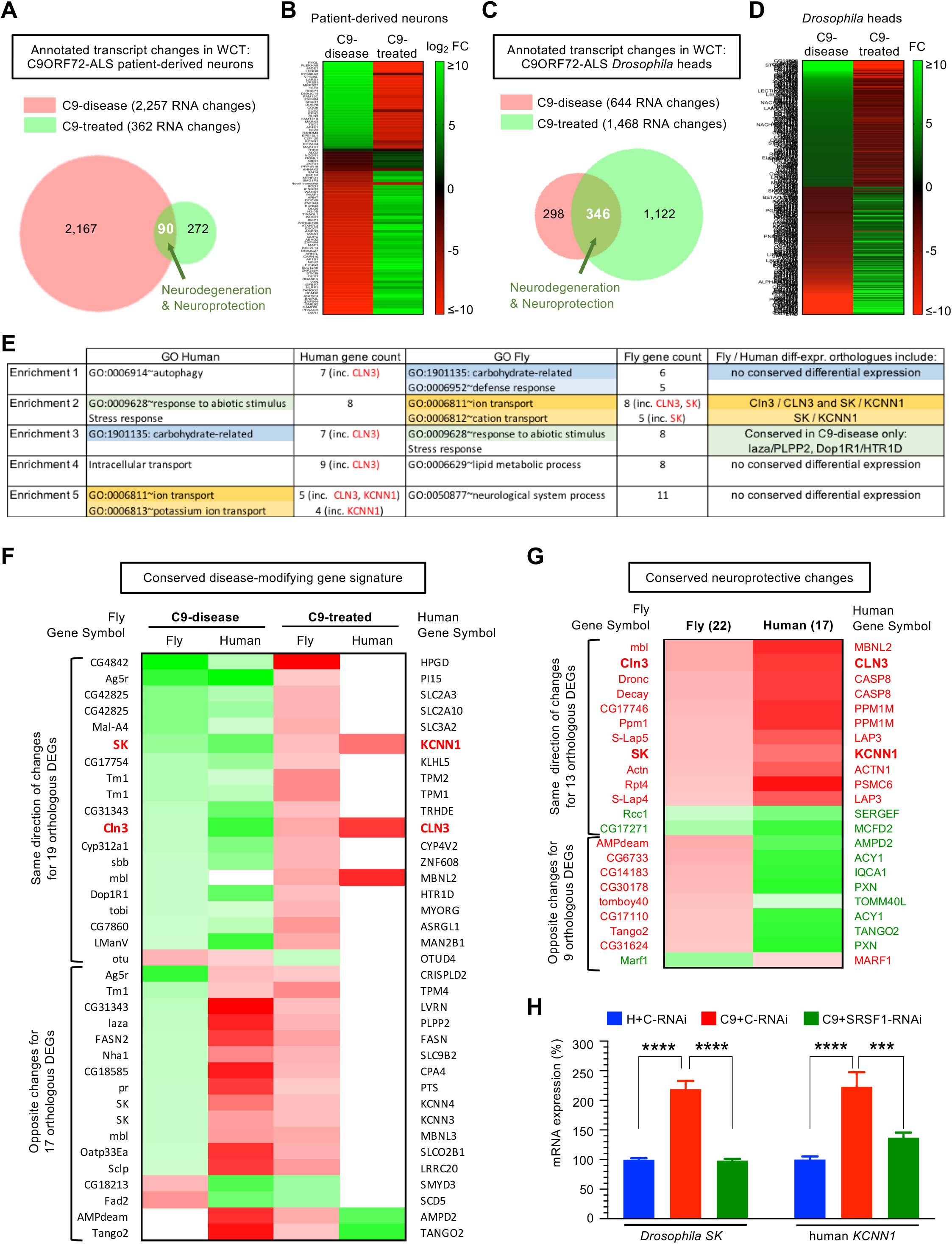
Identifying conserved disease-modifying gene expression signatures and neuroprotective transcripts. (**A**) Venn diagram representing differentially expressed genes (DEGs) identified in the human C9-disease and C9-treated groups. (**B**) Identification of a human disease-modifying gene expression signature. Heatmap representing the computed fold changes (FC) for the common human transcripts modulated in both C9-disease and C9-treated groups. Red labels show down-regulated transcripts while green depicts upregulated transcripts. (**C**) Venn diagram representing differentially expressed genes (DEGs) identified in the *Drosophila* C9-disease and C9-treated groups. (**D**) Identification of a *Drosophila* disease-modifying gene expression signature. Heatmap representing the FC for the common *Drosophila* transcripts modulated in both C9-disease and C9-treated groups. Red labels show down-regulated transcripts while green depicts upregulated transcripts. (**E**) Gene ontology analysis of conserved differentially-expressed transcripts identified in the human and *Drosophila* disease-modifying gene expression signatures carbohydrate metabolism, ion transport and response to stress as commonly altered in disease and manipulated upon neuroprotection. (**F**) Heatmap representing the computed fold changes for the orthologous genes in the disease-modifying signatures. Red labels show down-regulated transcripts while green depicts upregulated transcripts. (**G**) Heatmap representing the computed fold neuroprotective changes for the orthologous genes in the C9-treated groups. Red labels show down-regulated transcripts while green depicts upregulated transcripts. (**H**) qRT-PCR quantification of *Drosophila SK* and human *KCNN1* orthologous transcripts. Relative expression levels of *SK* mRNA was quantified for the indicated *Drosophila* lines in biological triplicates following normalization to Tub84b mRNA levels and to 100% for G4C2×3 + C-RNAi *Drosophila* heads (mean ± SEM; one-way ANOVA with Tukey’s correction for multiple comparisons; ****: p<0.0001; N (qRT-PCR reactions)=3). Relative expression levels of *KCNN1* mRNA was quantified in whole-cell patient-derived neurons in biological triplicates following normalization to U1 snRNA levels and to 100% for healthy neurons treated with C-RNAi (mean ± SEM; one-way ANOVA with Tukey’s correction for multiple comparisons, ***: p<0.001, ****: p<0.0001; N (qRT-PCR reactions)=3).

We further used BioMart [42] to identify conserved gene changes in the human and fly disease-modifying signatures. 40 differentially-expressed human genes out of 90 C9-disease/treated transcript changes have 49 fly homologues while 99 differentially-expressed fly genes out of 346 C9-disease/treated transcript changes have 78 human homologues (**Table S12 tabs 3 and 4**). Only 2 orthologous gene changes are commonly predicted to be up-regulated in the human and fly disease groups while their expression levels are down-regulated upon neuroprotection in both the human and fly C9-treated groups: human small conductance calcium-activated potassium channel protein 1 (*KCNN1*)/ fly small conductance calcium-activated potassium channel protein (*SK*) and the human and fly lysosomal/endosomal transmembrane proteins *CLN3/Cln3*. Gene ontology analysis was further performed on the 40 and 99 differentially-expressed and conserved genes respectively identified in the *in vitro* and *in vivo* disease-modifying gene expression signatures (**Table S12 tabs 5 and 6**). Top enriched categories in both human and fly include response to abiotic stimulus/ stress response, carbohydrate metabolism and cation/potassium transport (Figure 5E) indicating that if only 2 orthologous gene changes are commonly differentially-expressed, neuroprotection is achieved through manipulation of different genes which are involved in the same biological processes. Interestingly, the fully conserved C9-disease/treated changes in *CLN3/Cln3* and *KCNN1/SK* expression levels are both involved in the ion transport pathway. A heatmap of conserved genes showing at least one differentially-expressed change in the C9-disease or the C9-treated groups highlights 19 similar and 17 opposite direction of orthologous changes (Figure 5F; **Table S12 tab 7**). Most orthologous changes are identified in the C9-disease groups indicating that neuroprotection largely involves manipulation of different genes in C9ORF72-ALS patient-derived neurons and *Drosophila*.

To further investigate mechanisms of pathogenesis and neuroprotection, we identified the conserved gene changes in the C9-disease and C9-treated transcriptomes. 812 human differentially-expressed genes out of 1,804 DEGs found in C9-disease (Table S5) have 1,251 fly homologues and 65 differentially-expressed fly orthologues (**Table S13 tab 1**). 201 differentially-expressed fly genes out of 644 DEGs (Table S10) have 143 human homologues and 47 differentially-expressed human orthologues (**Table S13 tab 2**). A heatmap of the conserved changes identified in C9-disease is presented in Supplementary Figure S5, showing that only a small proportion of disease-altered gene expression changes have differentially-expressed orthologues. In addition to the challenge posed by integrating transcriptomes from human-derived neurons and *Drosophila* heads which contain multiple cell types, this might also be explained by the fact that only 17.8% and 24.4% of genes have annotated orthologues in the *Homo sapiens* and *Drosophila melanogaster Ensembl* genomes respectively (Supplementary Figure S6A-B). Likewise, 188 human differentially-expressed genes out of 351 neuroprotective gene changes have 261 fly homologues and 22 differentially-expressed fly orthologues (**Table S13 tab 3**) while 510 differentially-expressed fly genes out of 1,468 neuroprotective DEGs have 953 human homologues and 17 differentially-expressed human orthologues (**Table S13 tab 4**). Only 17 human and 22 fly orthologous genes are involved in conferring the SRSF1-RNAi dependent neuroprotection with 13 having similar changes of direction (Figure 5G). Conserved manipulation of these transcripts suggests that manipulation of other cellular pathways including various metabolic enzymes and apoptosis likely play important neuroprotective roles beyond modulation of *KCNN1/SK*, *CLN3/Cln3* and the ion transport pathway.

Alterations of voltage-gated ion channels have been reported in C9ORF72-ALS [54–57] and previous reports including patents for pharmacological modulation of SK channels in ALS indicate a potential role of small conductance Ca^2+^-activated potassium (SK) channels in ALS. We therefore experimentally validated our RNA-seq data using qRT-PCR assays to show that *KCNN1/SK* transcripts were indeed up-regulated in the C9-disease groups and down-regulated to normal expression levels upon SRSF1-RNAi-induced neuroprotection in the C9-treated groups (Figure 5H).

### Manipulating SK/KCCN ion channel activity alleviates C9ORF72-ALS motor neuron death and *Drosophila* locomotor deficits

We next wanted to functionally validate that our gene expression signature is enriched in genes with disease-modifying potential by testing the effects of manipulating the conserved *Drosophila* SK and human SK channel subunits in C9ORF72-ALS patient-derived neurons and *Drosophila*.

Four KCNN1-4 SK channel isoforms are found in the human genome. Using qRT-PCR quantification, we observed that *KCCN1* and *KCNN3* transcripts are significantly up-regulated in C9ORF72-ALS models while they are down-regulated upon neuroprotection (Supplementary Figure S7). On the other hand, *KCNN4* is not affected by the depletion of SRSF1, but is drastically reduced at mRNA level in C9ORF72-ALS patient-derived neurons. We next sought to functionally investigate the effects of inhibiting the up-regulated KCNN1 (encoding Kca2.1) and KCNN3 (encoding Kca2.3) transcripts in C9ORF72-ALS motor neurons. Motor neurons were generated from 2 lines each of healthy control (MIFF1, CS14) and C9ORF72-ALS patients (ALS-28, ALS-29; **Table 1**). Motor neurons were differentiated from neural progenitor cells, which express the characteristic Nestin and Pax6 markers (Supplementary Figure S8), through a 40-day differentiation protocol that leads to mature motor neurons expressing ChAT (Supplementary Figure S9). We further validated that C9ORF72-ALS motor neurons recapitulate RNA foci, a characteristic pathological hallmark of disease (Supplementary Figure S10). Interestingly, compared to healthy controls, C9ORF72-ALS motor neurons displayed significantly higher levels of caspase-3 positive cells, a marker of apoptosis (Figure 6A, top right bar chart), which correlated with a significant increase in nuclear fragmentation and apoptosis (Figure 6A, bottom right bar chart). This offers a unique opportunity to test the effects of the pharmacological inhibition of KCCN1/3 channels on C9ORF72-ALS motor neuron survival. The addition of increasing concentrations of apamin, an antagonist of KCNN1 and KCNN3 channels [34], to the motor neuron cultures resulted in a dose-dependent reduction of caspase-3 positive cells specific to C9ORF72-ALS motor neurons (Figure 6B). These data show that, consistent with the RNA-seq investigation, C9ORF72-ALS motor neurons have an enhanced expression of KCNN subunits and that inhibiting the activity of these Ca^2+^-activated potassium channels, as in the SRSF1-RNAi intervention, decreases apoptosis and cell death, promoting in turn neuronal survival.

**Fig. 6.**
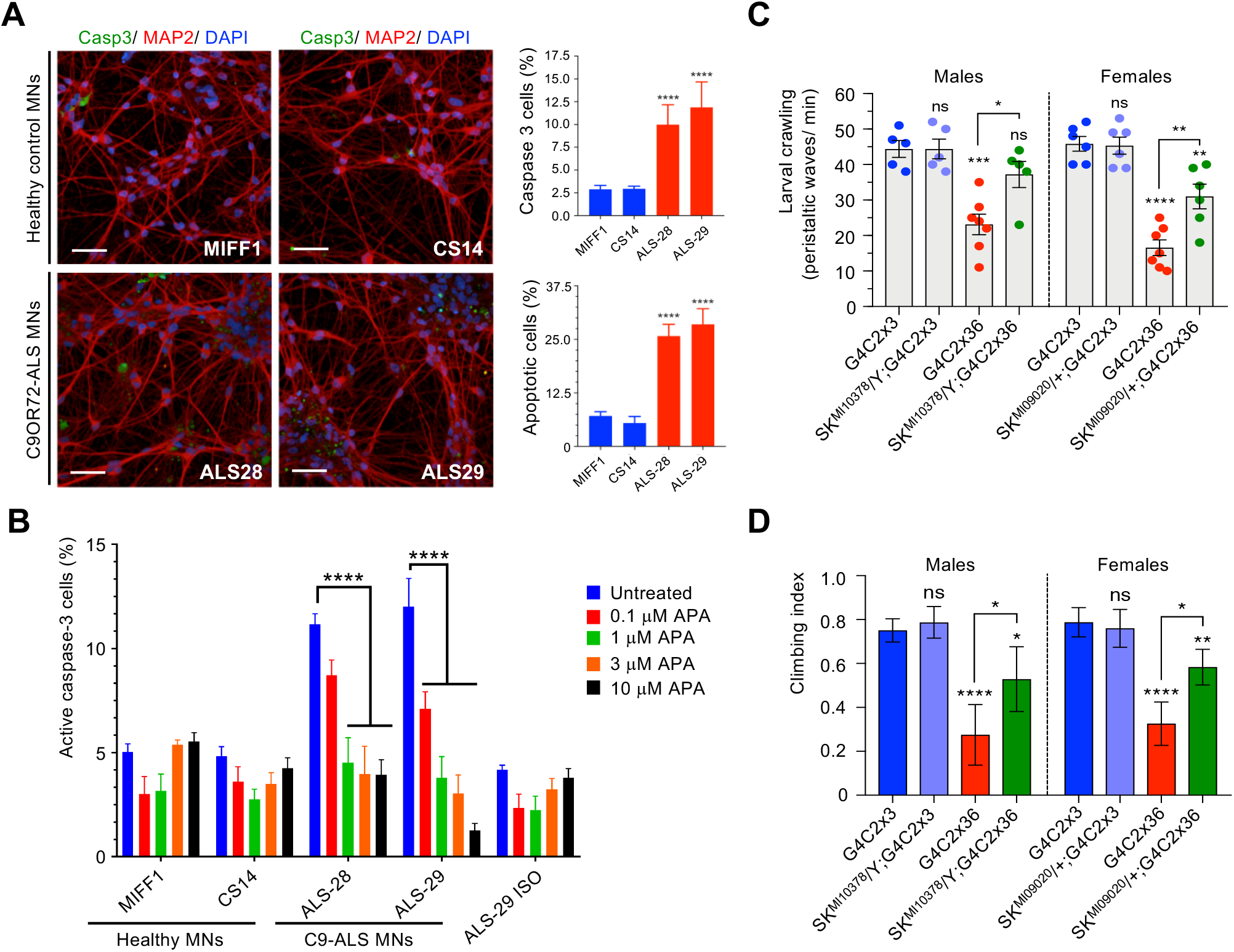
Pharmacological and genetic manipulation of KCCN channels promotes C9-ALS MNs survival and *Drosophila* locomotor function. (**A**) High content images of caspase 3 and MAP2 positive motor neurons from two control (MIFF1 & CS14) and two C9ORF&2-ALS (ALS-28 & ALS-29) lines. Scale bars: 50 µm. Bar charts represent the percentage of caspase 3 positive cells (top right) and of apoptotic cells measured by nuclear fragmentation (bottom right) in healthy and C9-ALS patient-derived motor neurons (mean ± SEM; one-way ANOVA Tukey’s multiple comparison test, ****: p<0.0001; N (replicate experiments)=3). (**B**) High-content microscopy automated imaging quantification of caspase-3 positive cells (%) treated with DMSO or increasing concentrations of the KCNN1/3 inhibitor apamin (APA) in healthy, C9-ALS and isogenic CRISPR-Cas9 mediated deletion of the repeat expansion (ISO) patient-derived motor neurons (mean ± SEM; two-way ANOVA Tukey’s multiple comparison test, ****: p<0.0001; N (replicate experiments)=3). (**C**) Larval crawling ability of male and female control (G4C3×3) or C9-ALS (G4C2×36) *Drosophila* crossed or not with two different *SK* mutant lines (mean ± SEM; one-way ANOVA; ns: non-significant, *: p<0.05, **: p<0.01, ****: p<0.0001; N (*Drosophila* larvae)>5). (**D**) Climbing ability of males and females control (G4C3×3) or C9-ALS (G4C2×36) *Drosophila* crossed or not with two different *SK* mutant lines (mean ± 95% CI; Kruskal-Wallis with Dunn’s correction; ns: non-significant, *: p<0.05, **: p<0.01, ****: p<0.0001; N (*Drosophila* flies)>25).

To further evaluate the neuroprotective potential of inhibiting the SK channel function in C9-*Drosophila*, Ctrl and C9 flies were crossed with two different loss-of-function *SK* mutant cell lines (Methods) and locomotor ability was analysed in hemizygous (male) or heterozygous (female) SK mutants. Strikingly, partial loss-of-function of SK channel activity significantly restored both the crawling activity of C9 larvae (Figure 6C) and the climbing ability of C9 adult flies (Figure 6D). Taken together, beyond highlighting the involvement of SK channels in C9ORF72-ALS, these data highlight the powerful approach of our study by targeting SRSF1-mediated transcriptomic changes to delineate the most C9ORF72-disease relevant mechanisms and neuroprotective strategies.

## Discussion

Multiple studies have reported thousands of transcriptome changes in C9ORF72-ALS neurons from cell and animal models as well as from post-mortem human brains, raising challenges for the identification of altered expression of transcripts that cause disease. Here, we combined for the first time *in vitro* and *in vivo* transcriptome investigations of C9ORF72-ALS human-derived neurons and *Drosophila* with a therapeutic disease-modifying strategy of neuroprotection which leads to specific inhibition of the SRSF1-dependent nuclear export of pathological *C9ORF72* repeat transcripts [12]. In perfect agreement with published transcriptome studies, we identified over 2,000 transcript changes in human C9ORF72-ALS affecting cellular pathways involved in neuronal-related processes, signalling, RNA metabolism, cell junction/ adhesion, cytoskeleton, cell death regulation and responses to stress [58–60]. Dysregulation in the expression of genes involved in synaptic-related processes, neuron differentiation and membrane hypoexcitability were also reported in C9ORF72-ALS human-derived neurons [58] while immune response and RNA processing alterations were also identified in a human post-mortem ALS brain study which investigated genome-wide expression changes using samples stratified by disease severity [61].

Strikingly, we discovered that manipulating the expression of a small proportion of transcripts dysregulated in human C9ORF72-ALS (362 transcripts genome-wide) is sufficient to confer therapeutic efficacy and neuroprotection. This also implies that the vast majority of diseased gene expression changes occur secondary to the neurodegenerative process and that a complete reversal of most altered gene expression changes is not required to achieve a neuroprotective effect. In both C9ORF72-ALS human-derived neurons and G4C2×36 *Drosophila*, the expression of approximately one third of the SRSF1-RNAi-manipulated transcripts altered in disease is completely reversed upon therapeutic depletion of SRSF1, revealing in turn *in vitro* and *in vivo* C9ORF72-ALS disease-modifying gene expression signatures. Identifying that of the thousands of RNA changes occurring in C9ORF72-ALS, manipulation of 261 human mRNAs, including 90 C9ORF72-ALS-modifying transcripts, is sufficient to confer neuroprotection provides a novel exciting basis to evaluate disease-modifying treatments, discover potential new biomarkers and importantly pinpoint a small number of new targets with therapeutic potential.

To demonstrate the power of our identification of disease-modifying transcript approach and emphasizing that our data integration leads to the identification of neuroprotective changes, we further showed that decreasing the activity of conserved SK potassium channels, which are upregulated in disease, mitigates the death of human-derived motor neurons as well as locomotor deficits in *Drosophila*.

Taken together, this investigation provides further validation that the partial depletion of SRSF1 is a promising gene therapy approach for neuroprotection, showing very limited effects in healthy or C9ORF72-ALS neurons at a global genome-wide level. Furthermore, minimal cellular splicing and nuclear export changes following SRSF1 depletion concur with the redundant roles of SRSF1-7 in the NXF1-dependent nuclear mRNA export adaptor function [25]. Consistent with this, simultaneous depletion of the general mRNA export adaptor Alyref and THOC5, two subunits of the Transcription-Export (TREX) complex which does not contain the SRSF1 protein [62, 63], leads to a drastic reduction in the recruitment of the nucleoporin-binding NXF1 export receptor onto polyadenylated mRNAs [64]. This indicates that TREX is responsible for the nuclear export of the bulk cellular mRNAs, a result recently confirmed by a comprehensive genome-wide investigation of the function of human TREX [65].

Finally, our data demonstrate genome-wide efficacy of the therapeutic depletion of SRSF1 with a remarkable mitigation of multiple unrelated cellular pathways altered in C9ORF72-ALS. In conclusion, the mechanisms of neuroprotection conferred by the partial depletion of SRSF1 involve both changes in the expression of a small number of neuroprotective transcripts encoding proteins involved in various functions (ion transport, RNA metabolism, synaptic transmission, etc.) as well as the inhibition of the nuclear export of *C9ORF72* repeat transcripts and the subsequent translation of dipeptide repeat proteins that also trigger gene expression alterations [66, 67]. The multiple additional neuroprotective benefits provided by the SRSF1-RNAi-induced manipulation of varied neuroprotective transcripts may explain why the partial suppression of one *ALYREF* gene, which was initially reported as a suppressor of *C9ORF72*-repeat mediated neurotoxicity in a *Drosophila* loss-of-function screen [68] and also identified as a binding partner of *C9ORF72*-repeat RNAs [35, 69], provide poor mitigating effects on the locomotor deficits of C9ORF72-ALS *Drosophila* [12]. Interestingly, altered cytoplasmic distribution of SRSF1 has been reported within the infarcted area of stroke victims when compared to the contralateral area, implicating a role for the cytoplasmic localisation of SRSF1 in tissue repair after ischaemia [70]. Several cytoplasmic related functions have also been reported for SRSF1, including nonsense-mediated decay (NMD) through the RS domain [26], mRNA degradation of specific targets [71], autoregulation of its mRNA through binding to its 3’UTR to prevent translation initiation [72], targeting of mRNAs to stress granules following stress [73, 74] and translation regulation [75–77]. It remains unknown whether this potential role in neuronal tissue repair is due to an impairment in the nuclear functions of SRSF1 or an enhancement of its cytoplasmic roles. Interestingly, our study which pinpoints neuroprotective benefits following partial depletion of SRSF1 would suggest that the nuclear reduction of SRSF1 may be the factor contributing to neuronal tissue repair after ischaemia through stimulation of the expression of transcripts involved in neuron differentiation, axonogenesis and synaptic transmission. Further investigation is now required to validate the potential safety and efficacy of the partial depletion of SRSF1 in the brain and spinal cords of pre-clinical mammalian models of C9ORF72-ALS/FTD.

## Conclusions

Thousands of gene expression changes, involving altered expression and splicing of transcripts, are typically identified in neurodegenerative diseases such as ALS. Here, we show for the first time that modulating 16% of human disease-altered transcripts only (362 out of 2,257 total pathological changes) is sufficient to confer the SRSF1-RNAi dependent neuroprotection previously identified as a promising novel gene therapy approach in C9ORF72-ALS patient-derived neurons and *Drosophila* [12]. Importantly, we found that the partial depletion of SRSF1 does not significantly alter transcriptomes (<1%) preserving the intrinsic variability of gene expression across individuals. It further leads to the therapeutic manipulation of key transcripts involved in the vast majority of C9ORF72-ALS-altered biological processes without significantly affecting genome-wide splicing (none) or mRNA nuclear export (0.4% transcriptome with modulations <2.5 fold). Overall, our genome-wide investigation provide a solid rationale for the therapeutic efficacy and safety of the partial depletion of SRSF1 *in vitro* and *in vivo* in preclinical patient-derived neurons and *Drosophila* models of C9ORF72-ALS. Integrating data between RNA-seq and microarrays as well as between diseased/neuroprotected human-derived neurons and *Drosophila* heads allowed identification of a disease-modifying signature with few but conserved neuroprotective targets with high therapeutic potential.

## Supporting information

Castelli et al. Supplementary Tables S1-13

## Abbreviations

ALS: amyotrophic lateral sclerosis
ANOVA: analysis of variance
C9ORF72: chromosome 9 open reading frame 72
DEGs: differentially-expressed transcripts at gene levels
KCNN: human small conductance calcium-activated potassium channel protein
MN: motor neuron
PCA: principal component analysis
SK: *Drosophila* small conductance calcium-activated potassium channel protein
SEM: standard error of the mean
SRSF1: serine/arginine-rich splicing factor 1

## Declarations

### Ethics approval and consent to participate

Informed consent was obtained from all patients before collection of fibroblasts under study STH16573 and Research Ethics Committee reference 12/YH/0330 (Prof Dame Pamela Shaw, University of Sheffield, UK). Induced pluripotent stem cell (iPSC) lines were obtained from Cedars-Sinai (USA), a nonprofit academic healthcare organisation. Patient-derived cell lines are described in Table 1.

## Acknowledgements

We acknowledge patients who donated fibroblasts which allowed the generation of patient-derived neurons. Micro-array analysis was undertaken by the Genomics Core Facility at the Sheffield Institute for Translational Neuroscience at the University of Sheffield. Data generation and processing of raw sequencing reads were carried out by the Centre for Genomic Research which is based at the University of Liverpool (project LIMS14705).

## Authors’ contributions

GMH and MM designed the transcriptomics studies. GMH, LF and AJW designed the experimental assays. PJS provided genetically characterised human biosamples. Computational analysis of the RNA-seq data was performed at gene expression level by LC and MM and at splicing level by IG and MRG. The *Drosophila* microarrays were run by PRH. LMC, CDSS and MAM performed the patient-derived neuron studies. ASM performed the *Drosophila* studies. GMH wrote the manuscript. LMC, PJS, AJW, LF and MM edited the manuscript. All authors approved the manuscript.

## Funding

This work was initially supported by the Medical Research Council (MRC) grant MR/M010864/1 (KN, GMH, PJS) and the MND Association grant Hautbergue/Apr16/846-791 (GMH, LF, AJW, PJS). This research is currently supported by the Biotechnology and Biological Sciences Research Council (BBSRC) grant BB/S005277/1 (GMH) and MRC New Investigator research grant MR/R024162/1 (GMH). LF was funded by the Thierry Latran Foundation (FTLAAP2016/ Astrocyte secretome) and is currently supported by the MND Association grant Apr16/848-791 and the Academy of Medical Sciences Springboard Award. CDSS is funded by an AstraZeneca Post-Doctoral award. AJW was supported by MRC core funding (MC_UU_00015/6) and ERC Starting grant (DYNAMITO; 309742). GMH also reports grants Apr17/854-791 from the MND Association, Thierry Latran FTLAAP2016/ Astrocyte secretome and Royal Society International Exchanges grant IEC\R3\17010 during the course of this study. MA acknowledge grants from Alzheimer’s Research UK (ARUK-PG2018B-005), European Research Council (ERC Advanced Award 294745) and MRC DPFS (129016). PJS is supported as an NIHR Senior Investigator Investigator (NF-SI-0617–10077), by the MND Association (AMBRoSIA 972-797), MRC grant MR/S004920/1 and the NIHR Sheffield Biomedical Research Centre: Translational Neuroscience for chronic neurological disorders (IS-BRC-1215-20017).

## Availability of data and materials

The RNA-seq and microarray data have respectively been deposited in *Gene Expression Omnibus* (GEO) under accession number GSE139900 (https://www.ncbi.nlm.nih.gov/geo/query/acc.cgi?acc=GSE139900) and GSE138592 (https://www.ncbi.nlm.nih.gov/geo/query/acc.cgi?acc=GSE138592). The R code for data analysis is available on request. All other data associated with this study are presented in the main text or Supplementary Materials.

## Competing interests

GMH, MA, AJW and PJS have USA-granted (US10/801027) and Europe-granted (EP3430143) patents for the use of inhibitors of SRSF1 to treat neurodegenerative disorders (WO2017207979A1).

The authors declare no other relationships, conditions or circumstances that present a potential conflict of interest.

## Supplementary Materials

**Supplementary Figures S1 to S10.**

**Table S1.** RNA-seq statistics

**Table S2.** Annotated quantified transcripts for RSEM>10

**Table S3.** Common cell marker counts

**Table S4.** SRSF1 transcript counts

**Table S5.** Differentially-expressed transcripts in patient-derived neurons

**Table S6.** Gene ontology analysis in patient-derived neurons

**Table S7.** Splicing analysis in patient-derived neurons

**Table S8.** Exon usage changes in patient-derived neurons

**Table S9.** mRNA nuclear export analysis in patient-derived neurons

**Table S10.** Differentially-expressed transcripts in *Drosophila*

**Table S11.** Gene ontology analysis in *Drosophila*

**Table S12.** C9ORF72-ALS disease-modifying gene expression signature

**Table S13.** Conserved human-fly gene expression changes

**Sequences of qPCR primers used in this study**

---

**Supplementary Figure S1.**
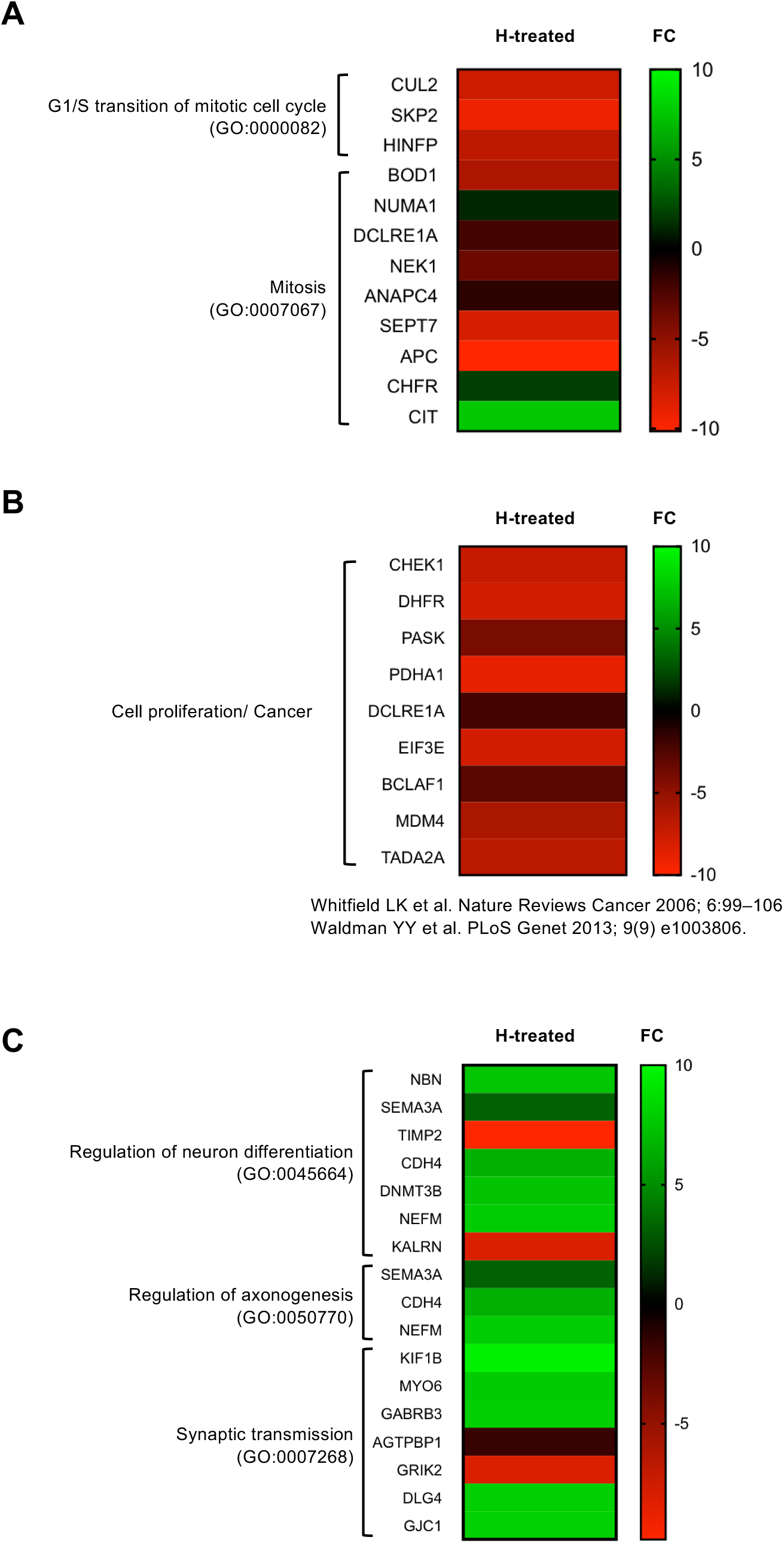
Downregulation of mitosis/ cell proliferation/ cancer makers and upregulation of neuronal-related functions in healthy neurons with partial depletion of SRSF1. (**A**) Heatmap representing the computed fold changes for the healthy-treated neurons group. Red labels down-regulated transcripts while green depicts upregulated transcripts. (**B**) Heatmap representing the computed fold changes for the healthy-treated neurons group. Red labels down-regulated transcripts while green depicts upregulated transcripts. (**C**) Heatmap representing the computed fold changes for the healthy-treated neurons group. Red labels down-regulated transcripts while green depicts upregulated transcripts.

**Supplementary Figure S2.**
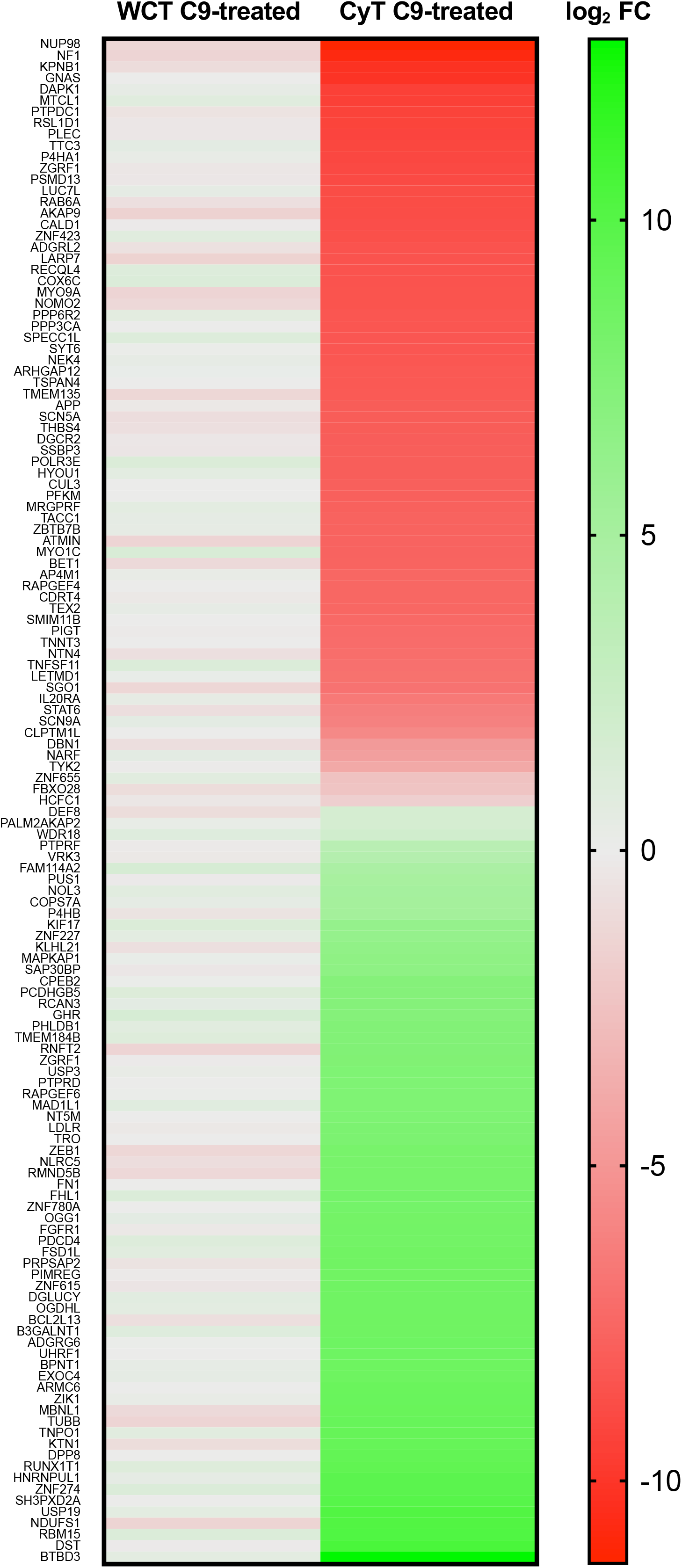
Enlargement of Fig. 3B. mRNA nuclear export analysis upon SRSF1 depletion at mRNA and FC levels.

**Supplementary Figure S3.**
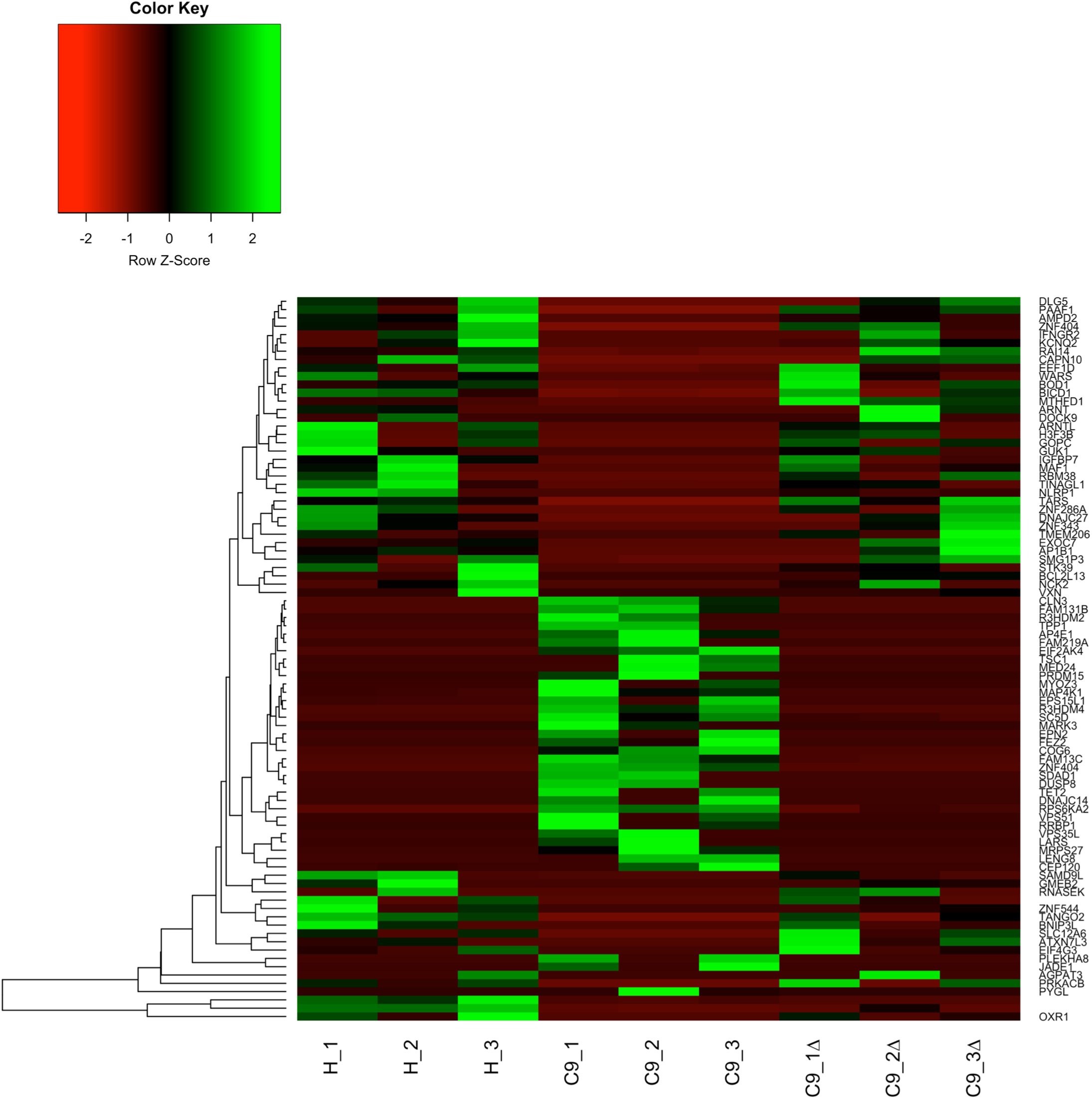
Heatmap representing a clustered analysis of the C9-ALS disease-modifying transcript signature in patient-derived neurons. H: healthy lines 1, 2, 3; C9: C9ORF72-ALS lines 1, 2, 3; Δ: SRSF1 depletion.

**Supplementary Figure S4.**
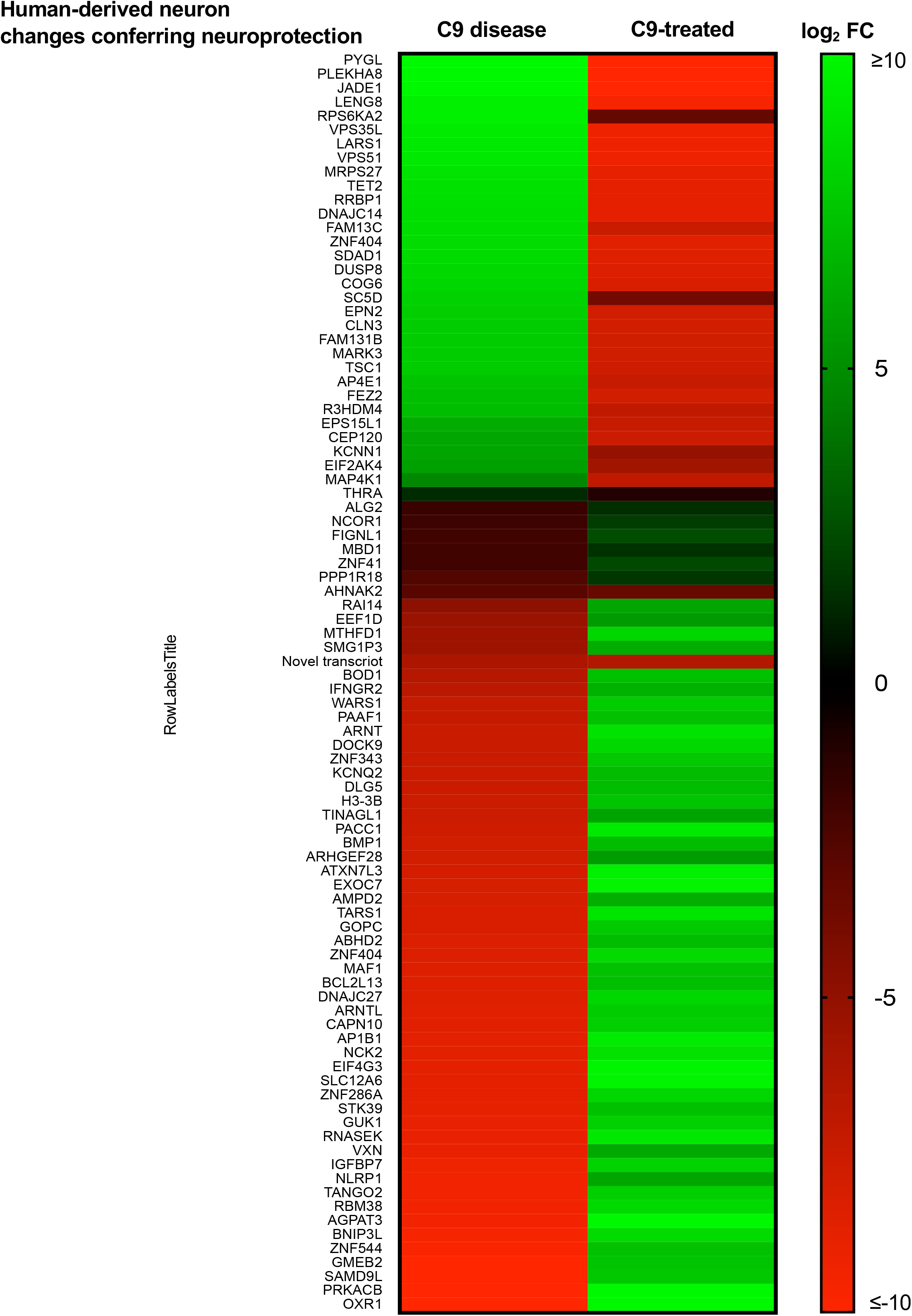
Enlargement of Figure 5B. C9-ALS disease-modifying signature at transcript ID and FC levels.

**Supplementary Figure S5.**
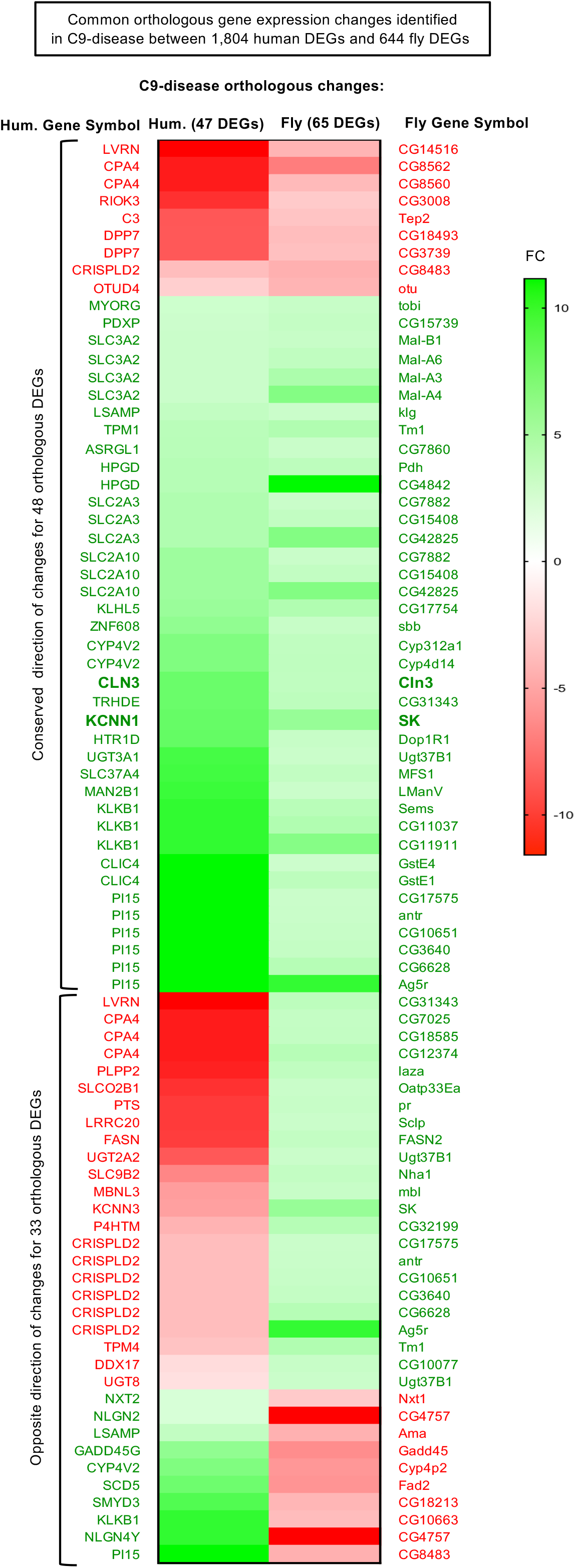
Orthologous gene changes in the human and fly C9-disease groups. 48 orthologues show conserved direction of differential expression at gene level (DEGs) while 33 others exhibit opposite direction of changes. Green and red labels respectively correspond to up-regulation and down-regulation of the expression levels of the corresponding transcripts.

**Supplementary Figure S6.**
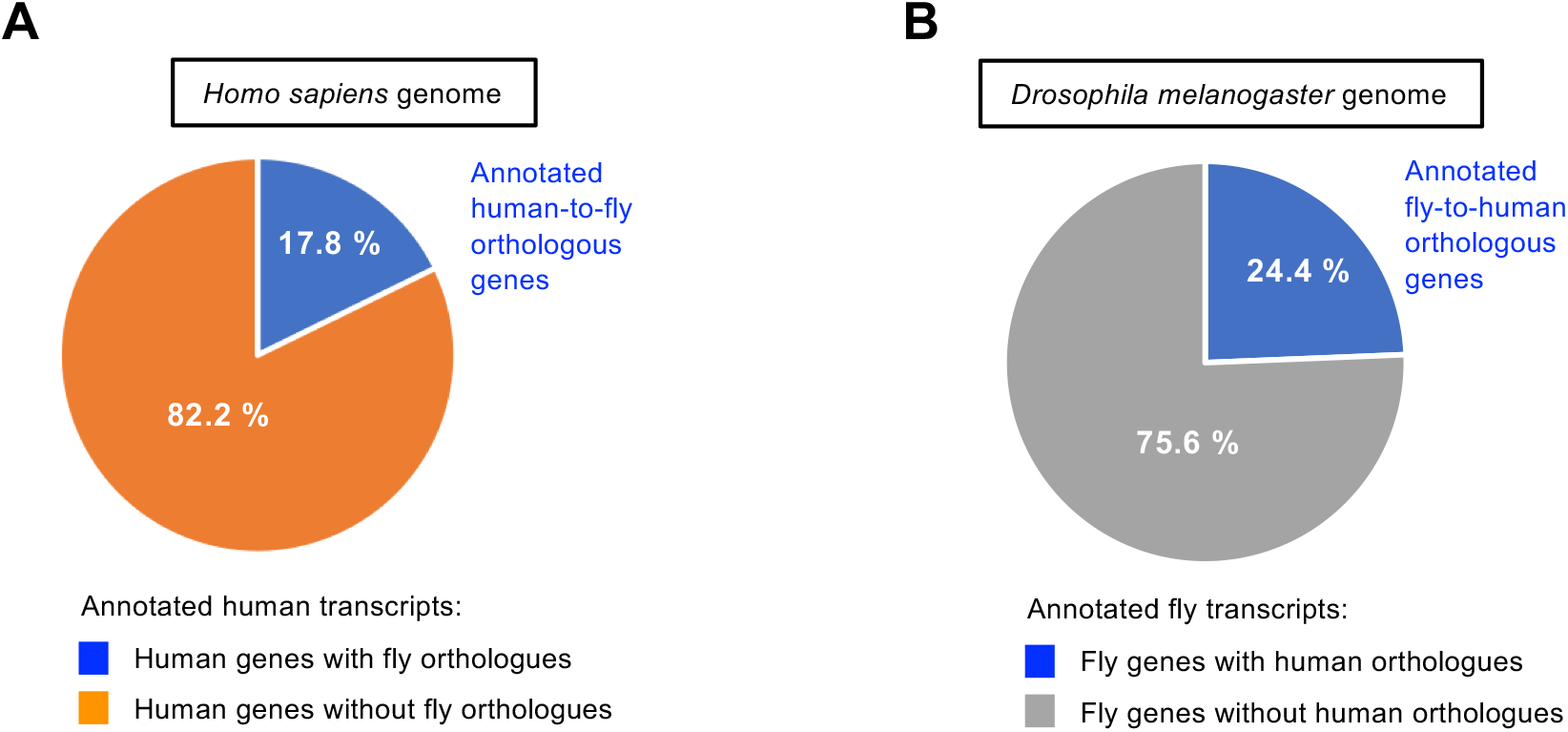
Orthologous genes in the human and fly genomes. BioMart (http://www.ensembl.org/biomart/martview) Ensembl Genes 99 was used with the *Homo sapiens* GRCh38.p13 and the *Drosophila melanogaster* BDGP6.28 genomes. A Multispecies comparison was performed on 25.04.2020 based on (i) Homologue filters, Orthologous genes only or (ii) Homologue filters, Orthologous human Genes only. (**A**) 3681 human coding genes out of 20,449 have conserved fly orthologous genes. (**B**) 3411 fly coding genes of 13,947 have conserved human orthologous genes. List of orthologous genes are available upon request.

**Supplementary Figure S7.**
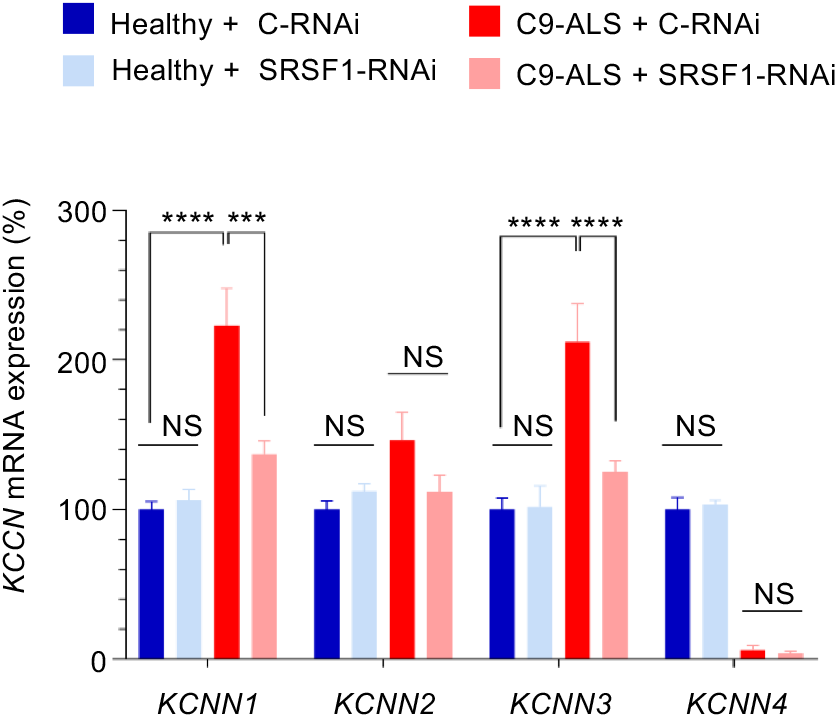
Quantification of KCNN1-4 mRNA expression levels in healthy and C9ORF72-ALS patient-derived neurons treated with control or SRSF1-RNAi lentivirus. Relative expression levels of *KCNN1*, *KCNN2*, *KCNN3* and *KCNN4* mRNAs were quantified using qRT-PCR in biological triplicates following normalization to *U1* snRNA levels and to 100% for healthy neurons treated with Ctrl-RNAi (mean ± SEM; two-way ANOVA with Tukey’s correction for multiple comparisons; NS: not significant; ***: p<0.001, ****: p<0.0001; N (qRT-PCR reactions)=3).

**Supplementary Figure S8.**
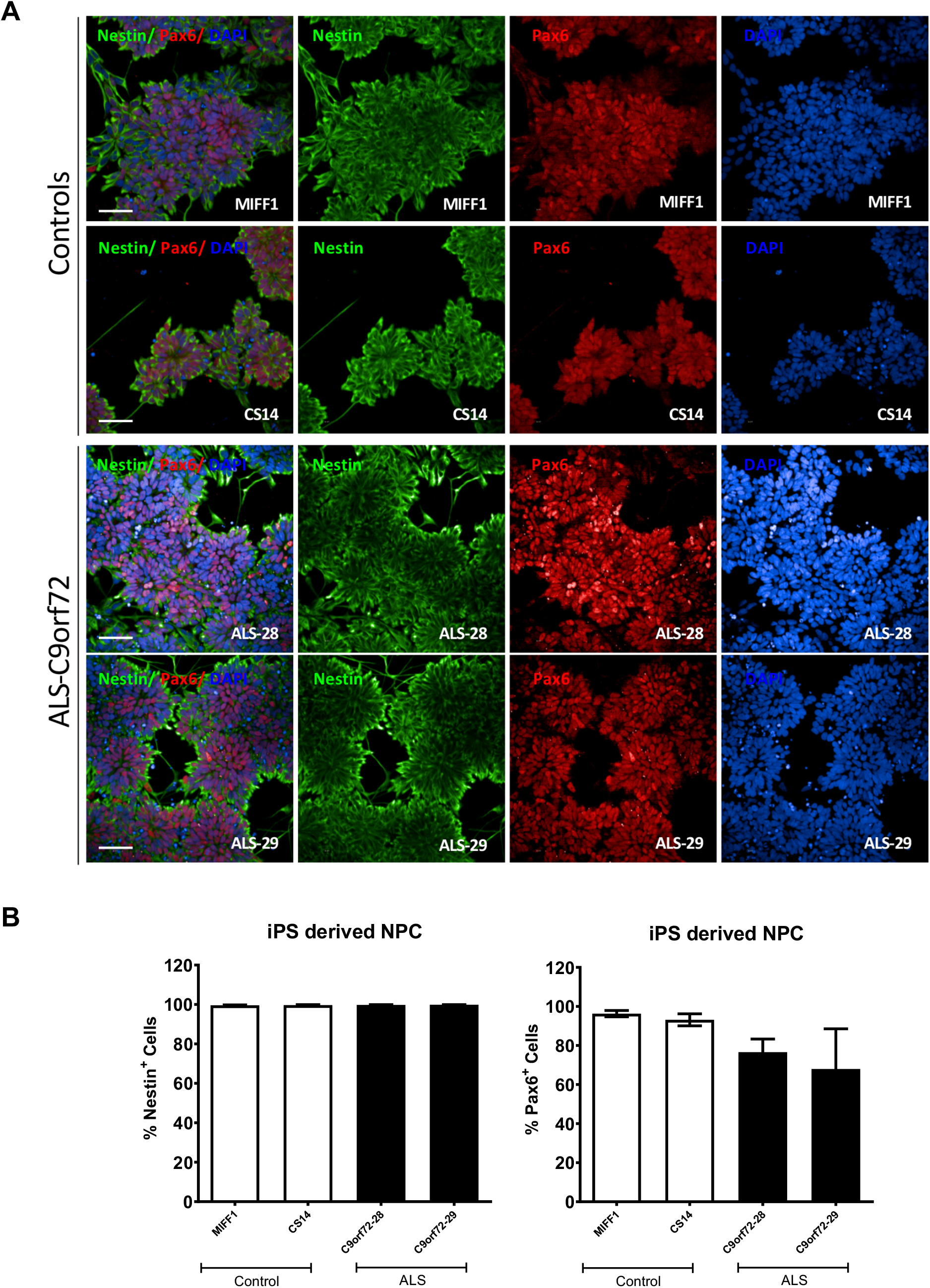
Characterisation of neural precursors cells (NPCs) derived from induced pluripotent stem cells (iPSCs) from healthy control and C9ORF72-ALS patients. (**A**) NPCs display positive staining for progenitor markers Pax6 and Nestin. Scale bars: 50 μm. (**B**) Quantification of cells positive for Nestin or Pax6 (n=3).

**Supplementary Figure S9.**
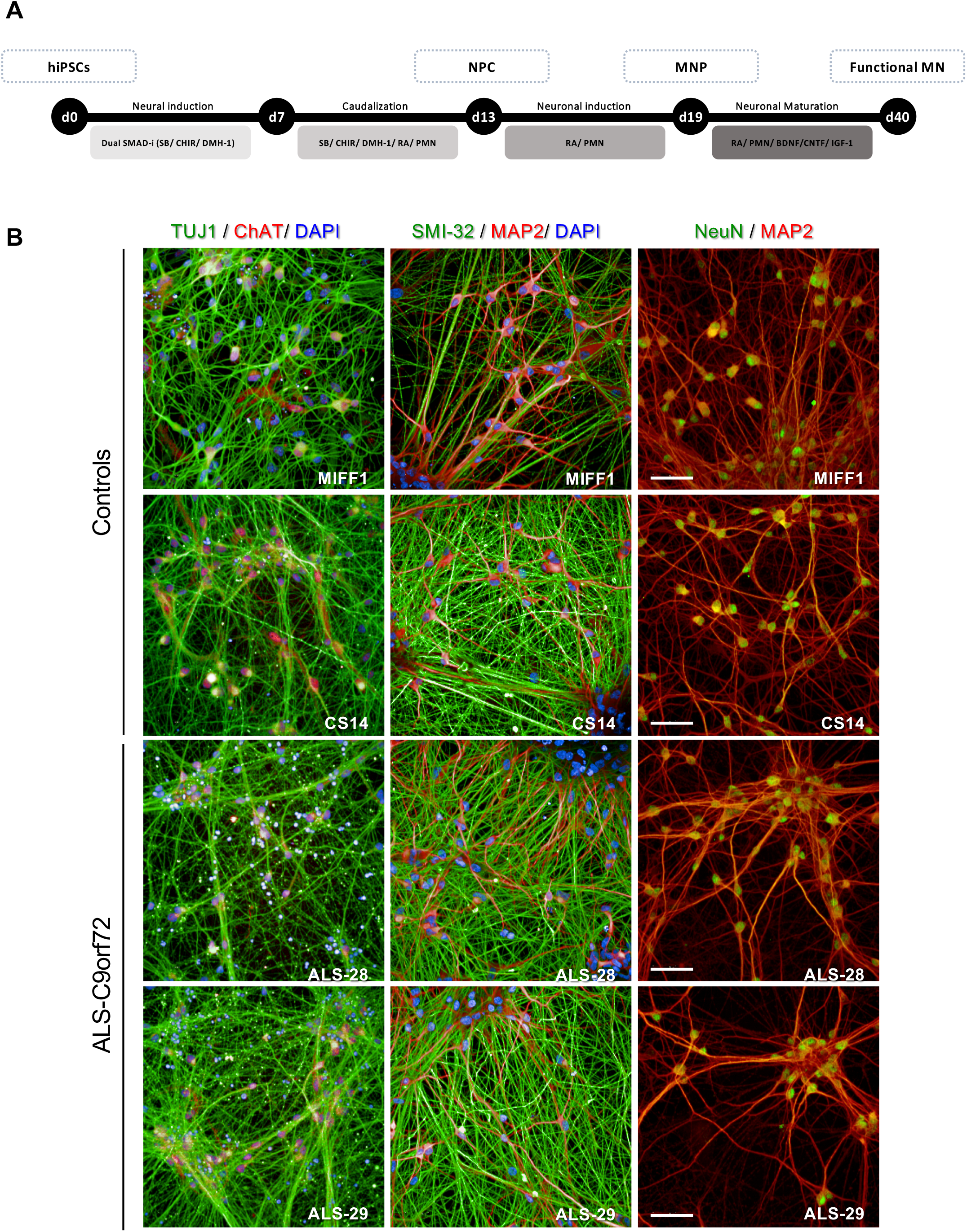
Differentiation of iPSCs from healthy and C9ORF72-ALS patients into functional motor neurons. (**A**) Schematic representation of the motor neuron differentiation protocol. (**B**) Representative images of motor neurons stained for the neuronal markers Tuj1, SMI-32, NeuN, MAP2 and the motor neuron specific marker ChAT. Scale bars: 50 μm.

**Supplementary Figure S10.**
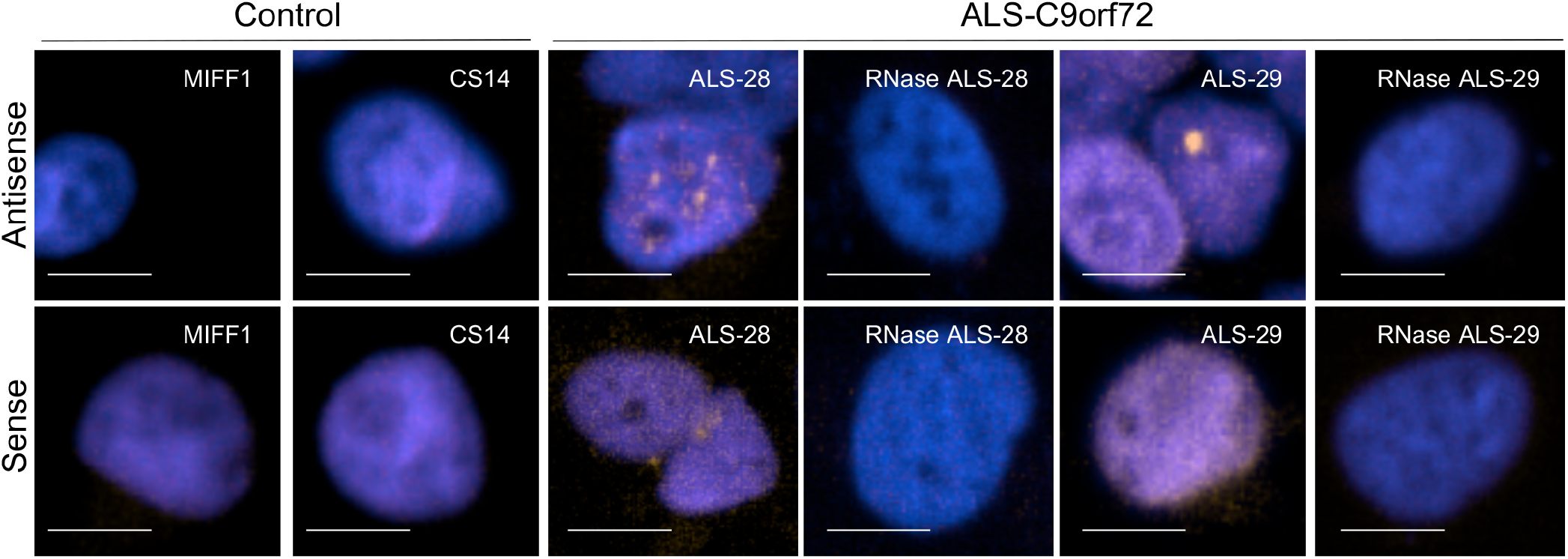
Motor neurons derived from C9ORF72-ALS patients develop characteristic sense and antisense RNA foci. Scale bars: 50 μm.

## Supplementary material: sequences of qPCR primers used in this study

Primers for *Drosophila Tub84b* (Ling *et al*., PLoS ONE 2011; 6:e17762):

Fwd: 5’-TGGGCCCGTCTGGACCACAA-3’

Rev: 5’-TCGCCGTCACCGGAGTCCAT-3’

Primers for *Drosophila SK* (designed using Primer-BLAST):

Fwd: 5’-ACCCTGTACTGCTGTTGCC-3’

Rev: 5’-TGTACAGATTCTGATGGATGGCTT-3’

Primers for *Drosophila NAAT1* (designed using Primer-BLAST):

Fwd: 5’-CACGGGATTGGCCTTCATCT-3’

Rev: 5’-CACGGGATTGGCCTTCATCT-3’

Primers for *Drosophila DHD* (designed using Primer-BLAST):

Fwd: 5’-GTGGTCCCTGCAAGGAAATG-3’

Rev: 5’-CACCTTGTAGCGCTCCGTC-3’

Primers for Human *U1 snRNA* (Hautbergue *et al*., 2017; 8:16063):

Fwd: 5’-CCATGATCACGAAGGTGGTT-3’

Rev: 5’-ATGCAGTCGAGTTTCCCACA-3’

Primers for Human *SRSF1* (Hautbergue *et al*., 2017; 8:16063):

Fwd: 5’-CCGCATCTACGTGGGTAACT-3’

Rev: 5’-TCGAACTCAACGAAGGCGAA-3

Primers for Human *C9ORF72* (Hautbergue *et al*., 2017; 8:16063):

Intron-1 Rev: 5’-GGAGAGAGGGTGGGAAAAAC-3’

Exon-3 Rev: 5’-GTCGACATGACTGCATTCCA-3’

Exon-1 For: 5’-TCAAACAGCGACAAGTTCCG-3’

Primers for Human *Usp49* (Origene):

Fwd: 5’-GGAGAATCTACGCTTGTGACCAG-3’

Rev: 5’-CGGAGAACCTGAGGTAGTCTGT-3’

Primers for Human *RSL1D1* (Origene):

Fwd: 5’-TCCGAAGACGAAATCCCACAGC-3’

Rev: 5’-GTGCTGGGATTAGGACTCTTTGC-3’

Primers for Human *MSH6* (Origene):

Fwd: 5’-AAGGACTGGCAGTCTGCTGTAG-3’

Rev: 5’-CGGCAACACAGAATTACTGGGCGA-3’

Primers for Human *RBM15* (Origene):

Fwd: 5’-CTTCCCACCTTGTGAGTTCTCC-3’

Rev: 5’-CTTCTTGTTCTCATACCTAACTCC-3’

Primers for Human *MTCL3* (Origene):

Fwd: 5’-TGCTCAAGTGCCGTCTGGAACA-3’

Rev: 5’-TGACTGTCTGCCAGGAGCTTCT-3’

Primers for Human *NUP98* (Origene):

Fwd: 5’-CCATCTATGGATGACCTGTAAA-3’

Rev: 5’-TCCGACCAATAGTGAAATCAGAGA-3’

Primers for Human *DAPK1* (Origene):

Fwd: 5’-CCAGACTGTCTTCCACCAACTC-3’

Rev: 5’-TCCTCACACTCACGTTCTCGCA-3’

Primers for Human *FN1* (Origene):

Fwd: 5’-GGACACAACGATGCTTCCTGAG-3’

Rev: 5’-ACAACACCGAGGTGACTGAGAC-3’

Primers for Human *USP19* (Origene):

Fwd: 5’-GCTGCTATCCTCAGAGTTGGCT-3’

Rev: 5’-TCATCCTCCGACTGTTGCTTCC-3’

Primers for Human *KCNN1* (Origene):

Fwd: 5’-TGCTGGTCTTCAGCATCTCCTC-3’

Rev: 5’-CGTAGCCAATGGAGAGGAAGGT-3’

Primers for Human *KCNN2* (Origene):

Fwd: 5’-GCCTTATCAGTCTCTCCACGATC-3’

Rev: 5’-CCAGTCATCTGCTCCATTGTCC-3’

Primers for Human *KCNN3* (Origene):

Fwd: 5’-GCCTTATCAGTCTGTCCACCATC-3’

Rev: 5’-TACAGGATGCGCTCGTAGGTCA-3’

Primers for Human *KCNN4* (Origene):

Fwd: 5’-CATTCCTGACCATCGGCTATGG-3’

Rev: 5’-GCCTTGTTAAACTCCAGCTTCCG-3’

Primers for Human *TRX1* (Origene):

Fwd: 5’-GTAGTTGACTTCTCAGCCACGTG-3’

Rev: 5’-CTGACAGTCATCCACATCTACTTC-3’

## Notes

### Competing Interest Statement

The authors have declared no competing interest.

