## Supplementary material for "Safety and efficacy of *C9ORF72*-repeat RNA nuclear export inhibition in amyotrophic lateral sclerosis": Castelli et al. Supplementary Tables S1-13: Table S8. Exon usage changes_ALS neurons.docx

**Table S8.** Statistically significant exon usage changes identified in WCT and CyT transcriptomes of human derived neurons treated with C-RNAi or SRSF1-RNAi lentivirus.

| Transcriptomes | Changes at exon level | Changes at gene level |
| --- | --- | --- |
| WCT: H_C-RNAi vs C9_C-RNAi (C9 disease) | 40 | 34 |
| WCT: C9_C-RNAi vs C9_ΔSRSF1 (C9-treated) | 0 | 0 |
| WCT: H_C-RNAi vs H_ΔSRSF1 (healthy-treated) | 68 | 61 |
| CyT: H_C-RNAi vs C9_C-RNAi (C9 disease) | 99 | 77 |
| CyT: C9_C-RNAi vs C9_ΔSRSF1 (C9-treated) | 0 | 0 |
| CyT: H_C-RNAi vs H_ΔSRSF1 (healthy-treated) | 6 | 6 |
